# A comparative study on the impact of five *Desmodium* species on soil microbiome reveals enrichment of selected bacterial and fungal taxa

**DOI:** 10.1101/2023.02.07.527423

**Authors:** Aneth Bella David, Kilaza Samson Mwaikono, Charles Midega, Francis Magingo, Beatrix W. Alsanius, Laurie E. Drinkwater, Teun Dekker, Sylvester Lyantagaye

## Abstract

**Introduction:** Several *Desmodium* spp. are used as intercrops in push-pull pest management systems to repel insect herbivores. In addition, *Desmodium* suppresses the parasitic weed *Striga*, and diversifies the soil microbiome with negative impacts on fungi. We investigated the impact of a 2-year cropping of five *Desmodium* species on soil microbiome populations.

**Methodology:** Total DNA was obtained from root zone soil samples collected from a two-years-old common garden experiment with replicated plots of five *Desmodium* spp. at the international centre for insect physiology and ecology (ICIPE), Mbita, Kenya. Subsequently, 16S and ITS DNA sequencing were performed and the data was analysed by using QIIME2 and Calypso.

**Results:** Our findings show significant differences in composition and abundance of specific microbial taxa among the *Desmodium* plots and the bulk soil, with a stronger shift observed for fungal community profiles than bacteria. There was, however, no significant difference in overall diversity, richness and evenness of microbial communities among the *Desmodium* plots and the bulk soil. Similarly, beta diversity analysis did not reveal a significant association of variation to specific *Desmodium* spp. plots.

**Discussion and conclusion:** This is the first study to compare impact and association of whole soil microbiomes to different *Desmodium* species. Whereas long-term *Desmodium* cropping clearly shifts whole microbiome communities, no significant difference in overall diversity and richness of microbial populations was observed among the studied plots. However, there was a divergence of individual taxa reflected on their increased abundance in association to specific *Desmodium* spp., pointing towards potential impact on ecosystem services. These findings indicate that significant shifts in whole microbial populations due to *Desmodium* spp. and thus potentially provision of associated ecosystem services require longer cultivation periods to solidify. Future studies should focus on techniques that monitor real-time changes in microbial populations such as RNA-seq to ascertain live and dead microbes, and thus infer ecological services.

## INTRODUCTION

Push-pull technology for management of Lepidopteran insect pests of cereals employs a stimulo-deterrent mechanism, where *Desmodium* spp. intercrops play a critical role. Smallholder farmers in Eastern and Southern Africa grow *Desmodium* spp. between rows of cereal crops such as maize and sorghum to lower populations of deleterious insect pests away from the main crop. At the same time, *Brachiaria* cv. Mulato or Napier grass (*Pennisetum purpureum*) is planted on the border of the fields to attract and trap insect pests (Pickett et al., 2014). The technology effectively and sustainably controls stem-borers (*Chilo partellus* and *Busseola fusca*) and recently fall armyworms (*Spodoptera frugiperda*). In addition, the root exudates of Desmodium spp. intercrops suppress the parasitic weed *Striga*, endemic in Eastern Africa, thus leading to increases in cereal yield from reduced attack of both insect pests and the weed (Midega et al., 2015, 2018).

*Desmodium* spp. is a genus of flowering plants in the Fabaceae family of about 350 species that grow mainly in tropical and subtropical zones worldwide. Species of this genus find numerous uses including in traditional medicine (Ma et al., 2011; Farid et al., 2018) and commonly used as animal fodder (Heuzé et al., 2015, 2017). More importantly, species within the *Desmodium* genus were selected for use in a push-pull farming system through careful studies by scientists in Kenya, delivering multiple benefits in smallholder cereal cropping systems. Being leguminous plants, cultivation of *Desmodium* spp. also improves soil fertility through a range of mechanisms including nitrogen fixation, which is enhanced by the perennial nature of *Desmodium* spp. In addition, the intercrop builds soil organic matter reserves and promotes aggregate formation both of which can improve soil moisture conservation (Drinkwater et al., 2021). These soil health benefits likely further contribute to increases in yield of cereal crops, making push-pull an attractive system for smallholder farmers.

The diverse aspects of plant-plant and insect plant interactions in push-pull technology have been well investigated and documented. However, belowground interactions, especially with focus to soil microorganisms have not been investigated, despite the numerous ecological services provided by push-pull technology, in addition to pest control, some with clear indications of the role of soil microorganisms. Taking into account belowground interactions between plants and soil communities is of paramount importance when selecting intercrops considering the contribution of both individual species as well as whole soil microbiomes on plant health and ecosystem functioning (Compant et al., 2019). Yet this area is poorly researched especially regarding soil microbial shifts caused by closely related species or cultivars.

The composition and diversity of soil microbes at a particular location is determined by both biotic and abiotic factors, with aboveground vegetation having the largest influence (Philippot et al., 2013). Plant-soil microbes interactions are mediated through root exudates that provide an important source of carbon for microorganisms as well as signalling compounds (Haichar et al., 2014). In turn, soil microorganisms and nematodes play key roles in maintenance of soil structure and function through provision of critical ecological services. For instance, soil microbes play key roles in decomposition of organic matter and cycling of nutrients, carbon sequestration, promotion of plant health through bio-protection (Jacoby et al., 2017; Saccá et al., 2017) with recent studies suggesting that plant-associated microbes including those in the soil are involved in regulating plant-insect herbivore interactions (Friman et al., 2021; Grunseich et al., 2020; Pangesti et al., 2015). These findings suggest direct implications for farming practices where insect pests continue to devastate productivity. Understanding how specific crop plants modulate overall soil microbiota, and not just individual species, may enhance existing benefits as well as unlock new avenues for sustainable plant health and productivity improvement.

In a previous study we characterised the difference in soil microbial composition, structure and diversity in long term push-pull plots compared to maize monoculture (Mwakilili et al., 2021). Several other studies reporting on benefits of push-pull technology that are clearly linked to soil microbial communities suggest a deeper role than currently known. For example, a study by Njeru et al. (2020) revealed that maize coming from push-pull plots had lower levels of mycotoxins and mycotoxin-producing fungi compared to that from monoculture. In a separate study, the frequency of occurrence of a mycotoxin-producing fungus *Aspergillus flavus* was lower in push-pull than maize monoculture plots (Maxwell et al., 2017). These findings are in line with our previous study where we show that push-pull farming and *Desmodium* intercropping in cereal farming impact diversity of fungal communities more than bacteria (Mwakilili et al., 2021), manipulation of fungal communities in the soil to promote competitive beneficial filamentous fungi has been established as a method to manage mycotoxins in cereals (Sarrocco et al., 2019). In a similar vein, another study demonstrated that maize growing on soil from long-term push-pull plots produced higher amounts of secondary metabolites and experienced lesser herbivory than maize growing on soil from corresponding maize monoculture plots (Mutyambai et al., 2019). Although not investigated, these observations point towards the role of *Desmodium* intercrops in shaping soil microbial communities in push-pull farms and the subsequent microbial activities in plant-soil feedback mechanisms. Although we have already shown that long-term push-pull farming (*Desmodium* spp. intercropping) cause significant shifts in composition and structure of soil microorganisms, however, the time-scale of such changes, and whether the impact on soil microbial communities may be different between *Desmodium* species due to potential differences in composition of root exudates, is unknown. Understanding the differences in soil microbial associations to *Desmodium* spp. may show inter-linkage to the health of *Desmodium* spp. and their abilities to survive under diverse environmental stresses.

In the current study, we investigated the impact of five *Desmodium* species on soil microbial profiles. Currently, two *Desmodium* spp. are commonly used in push-full farming as intercrops, *D. intortum* (greenleaf desmodium) and *D. uncinatum* (silverleaf desmodium). These species have demonstrated several challenges including sensitivity to drought (*D. intortum*) and difficulty of producing seeds (personal communication). In search of more resilient *Desmodium* varieties suited for the varying African climates, several *Desmodium* species accessions were compared for their ability to withstand abiotic stresses including drought tolerance, these are *D. incanum, D. repandum, D. uncinatum, D. intortum* and *D. ramosissimum*, where *D. incanum* and *D. ramosissimum* were shown to exhibit stronger drought tolerance than the other *Desmodium* species, as well as stronger capability to suppress *Striga* weeds (Midega et al., 2017). We thus aimed at complementing the selection of the *Desmodium* spp. as intercrops in cereal push-pull farming by providing insights on their impact on soil microbial populations.

In the present study we show the composition of soil microbial communities in plots cultivated with five different *Desmodium* spp. in comparison to the bulk soil. We also highlight diversity measures as well as enriched taxa associated with each *Desmodium* spp. and the bulk soil.

## METHODOLOGY

### Sampling site

Soil samples were collected from ongoing common garden experimental plots at the International Centre for Insect Physiology and Ecology (ICIPE), Mbita campus, Kenya (0°25′S, 34°12′E). Mbita is located on the eastern shores of Lake Victoria, 1125 m above sea level. The area receives about 1001 mm rainfall per year and has an average annual temperature of 22.6 °C. Sampling was done during the cool dry season in July 2017. The soil type of the area is sandy loam/black cotton soils.

### Soil samples

Soil samples were collected from the common garden plots in which five different species of *Desmodium* had been growing for two years in 7.8 m^2^ plots in a completely randomised design. The five species have been under evaluation for use in push-pull systems in different agro-ecological regions of Kenya and included *Desmodium* spp.: *D. ramosissimum, D. repandum, D. uncinatum, D. intortum* and *D. incanum*. All plots were treated equally with no additives throughout the cultivation period. The plots relied on seasonal rainfalls and irrigation during the dry season. A 2 m buffer strip of bare soil from which control bulk soil samples were collected, separated the plots from the surrounding uncultivated grass-covered land.

For each treatment, three samples were collected. Three plots were selected from each *Desmodium* spp. treatment, with each plot representing one sample. Each sample was made up of a composite of three 15 – 18 cm deep cores taken randomly across *Desmodium* plots close to the roots (root zone). A total of three bulk soil control samples were also collected from the buffer zone where plants were constantly removed so that bare soil was left. Here also each sample was made up of a composite of three 15 – 18 cm deep cores. Afterwards, the composite soil samples from each plot and the buffer zone were homogenised and sieved through a 4 mm wire mesh. About 200 g soil sub-sample was then collected and stored at -20 °C for further analysis.

### DNA extraction and sequencing

DNeasy Powersoil kit (Qiagen, Manchester, UK) was used for total DNA extraction from the soil samples following the manufacturer’s protocol. Nanodrop spectrophotometer and gel electrophoresis were used to assess the quality, size and quantity of the extracted DNA. DNA samples were stored at -20 °C.

For bacterial communities, the V1-V3 region of the 16S rDNA gene was targeted with primer pairs 27F and 518R while ITS1F and ITS2 primer pairs were used for fungi targeting the ITS1 region.

Resulting amplicons were gel purified, end repaired and illumina specific adapter sequence were ligated to each amplicon (NEBNext Ultra II DNA library prep kit). Following quantification, the samples were individually indexed (NEBNext Multiplex Oligos for illumina Dual Index Primers Set 1), and another AMPure XP bead based purification step was performed. Amplicons DNA sequencing was done at Inqaba Biotechnical Industries (Pty) Ltd (Pretoria, South Africa) on Illumina MiSeq platform using a MiSeq v3 kit with 600 cycles (300 cycles for each paired read and 12 cycles for the barcode sequence) according to the manufacturer’s instructions. Demultiplexed 300bp paired-end reads were obtained.

### Bioinformatics and statistical analysis

FASTQC (Wingett & Andrews, 2018) was used to assess the quality of raw sequence reads. The reads were then imported into QIIME2 v2020.11 (Bolyen et al., 2019) where quality control, construction of a feature table and taxonomic classification were performed. In summary, quality control was done by using the dada2 plugin (Callahan et al., 2016) by trimming and truncating both the 16S and ITS reads to remove low quality parts. Taxonomic assignment was done by using feature-classifier classify-sklearn (Bokulich et al., 2018) by using pre-trained classifiers. Bacterial taxonomic assignment was based on Greengenes reference database (DeSantis et al., 2006) pre-trained on V1-V3 region of the 16S, while for the fungi, the UNITE v8.2 reference database (Nilsson et al., 2018) pre-trained to ITS1 was used. Important commands and parameters used are highlighted in Table 1.

**Table 1:** Commands and parameters used during data analysis in Qiime2

| Function | Command and parameters | Platform |
| --- | --- | --- |
| Trimming 16S sequences | --p-trim-left -f 8<br>--p-trim-left -r 8 | qiime2 |
| truncation 16S sequences | --p-trunc-len -f 290<br>--p-trunc-len -r 260 | qiime2 |
| Trimming ITS sequences | --p-trim-left 10 | qiime2 |
| Truncation ITS sequences | --p-trunc-len 299 | qiime2 |

Further, the feature table was converted into biom format (using qiime 2 export tool), and then imported into calypso V8.84 (http://cgenome.net:8080/calypso-8.84) (Zakrzewski et al., 2017) where further statistical and diversity analyses were performed. Before the analyses in calypso, samples with less than 1000 sequence reads, taxa with less than 0.01% relative abundance and taxa with over 50% zeroes were filtered out. Feature reads counts were normalized by total sum of squares (TSS) and transformed by both cumulative sum-scaling (CSS) and log2 to account for the non-normal distribution of taxonomic counts.

In calypso, different quantitative measures were analysed and plotted including taxa abundance and differential abundances in the treatments. Bray-Curtis distance metric was used to perform multivariate statistical testing and generate relevant plots for beta diversity estimation among the *Desmodium* spp. and control plots. Alpha diversity measures Shannon index, richness and evenness were also calculated as well as differential abundance and group association analyses.

## RESULTS AND DISCUSSION

In this study, we hypothesised that continuous cultivation of *Desmodium* species for 2 years caused shifts in soil root zone microbial community structure, composition and diversity relative to the bulk soil. With aboveground vegetation having been shown to exert the biggest influence on composition and structure of soil microbial populations (Hooper et al., 2010, 2015) we further hypothesised that the impact on soil microbial communities would diverge between *Desmodium* species due to potential variation in composition of root exudates. It was expected that the different species *Desmodium* would attract different assemblages of soil microorganisms in the root zone, with potential implications on health and functioning of *Desmodium* and the ecosystems of which they are part, such as in push-pull farming.

The findings show different aspects of soil microbial communities associated with the studied *Desmodium* species in contrast to the bulk soil; 1) differences in composition and abundance of soil microbes between *Desmodium* plots and bulk soil 2) highlight dominant soil microbial taxa associated with *Desmodium* spp., 3) unique and common microbial groups associated with *Desmodium* spp. as well as 4) diversity measures of soil microbial communities.

### Composition and abundance of soil microorganisms

A total of 15 bacterial and 8 fungal phyla were identified in all soil samples. The most abundant bacterial phyla were Chloriflexi (23%), Actinobacteria (21%) Cyanobacteria (15%), Acidobacteria (14%), Proteobacteria (8%) and Planctomycetes (8%). Other phyla included Bactroidetes, Gammatimonadetes, Nitrospirae, Elusimicrobia, Firmicutes and Armatimonadetes while two phyla were unclassified. Relative abundances of the identified bacterial phyla are shown in figure 1A.

**Figure 1:**
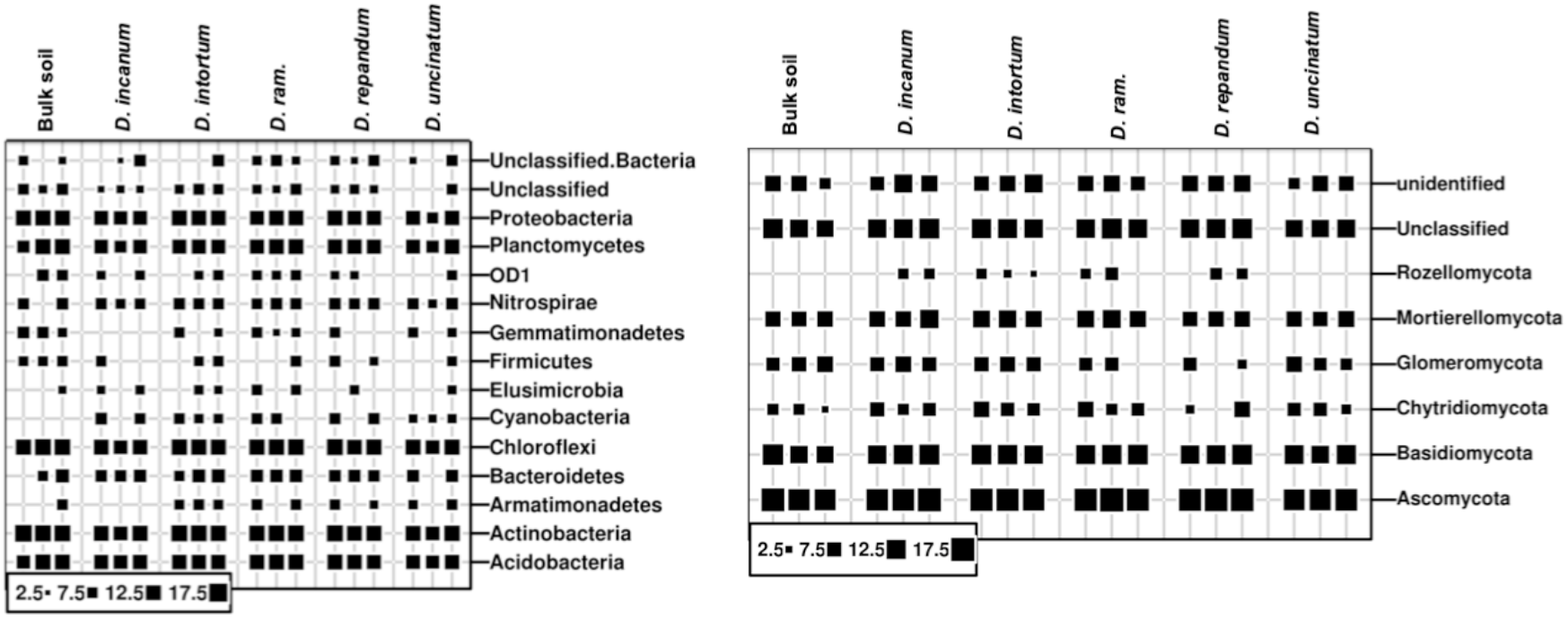
Bubble plots showing relative abundances of bacterial (left) and fungal (right) phyla in *Desmodium* spp. root zone soil and bulk soil. The relative abundances are shown as bubbles, with the size of the bubble being directly proportional to the relative abundance. Relative abundance was calculated from read counts (ASVs) normalized by total sum of squares (TSS) and transformed by cumulative sum scaling (CSS).

The majority of the fungal microbes belonged to the phylum Ascomycota (84%), followed by Basidiomycota (7.6%). Other phyla included Chytridiomycota (0.16 %), Glomeromycota (0.9 %) and Mortierellomycota (0.9 %). One phylum was unclassified and another unidentified. The relative abundances of fungal phyla in all plots are shown in Figure 1B.

At the genus level, most observed bacterial genera were unclassified due to the limitations of the classification databases in addition to potentially novel soil bacteria genera that may have not been classified in the past. However, among the few that were identified *i*.*e*., *Rhodoplanes, Gemmata, Nitrospira, Bradyrhizobium, Balneimonas, Streptomyces* and *Steroidobacter* occurred in varying abundances in all *Desmodium* spp. plots and bulk soil. The abundances of the 30 most abundant bacterial genera including those mentioned above are shown in figure 2 Compared to bacterial taxa, the majority of the abundant fungal genera were classified, as shown in figure 3, allowing for theorization of function based on literature. Both the *Desmodium* spp. plots and bulk soil harboured a diverse number of genera in varying abundances, with the genus *Fusarium* being the most Studies on shifts of soil microbial populations as a result of *Desmodium* spp. cultivation are scant. Literature on soil microbes and *Desmodium* spp. is populated by research on endophytes and nodule symbionts, in particular bacterial endosymbionts. Endosymbionts and nodule bacteria of other leguminous plants have been widely characterised and studied for their role in nitrogen fixing, an important ecological function. For *Desmodium* spp. (Parker, 2002) isolated several *Bradyrhizobia* species from *D. grahamii* nodules while (Toniutti et al., 2017) did the same from *D. incanum*. Most of the endosymbionts isolated from different *Desmodium* species in these studies tend to fall under three rhizobia genera; abundant taxa in both *Desmodium* species plots and the bulk soil. Other abundant genera identified are *Didymella, Chaetomium, Cladorrhinum, Stachybotrys* and *Curvularia*.

**Figure 2:**
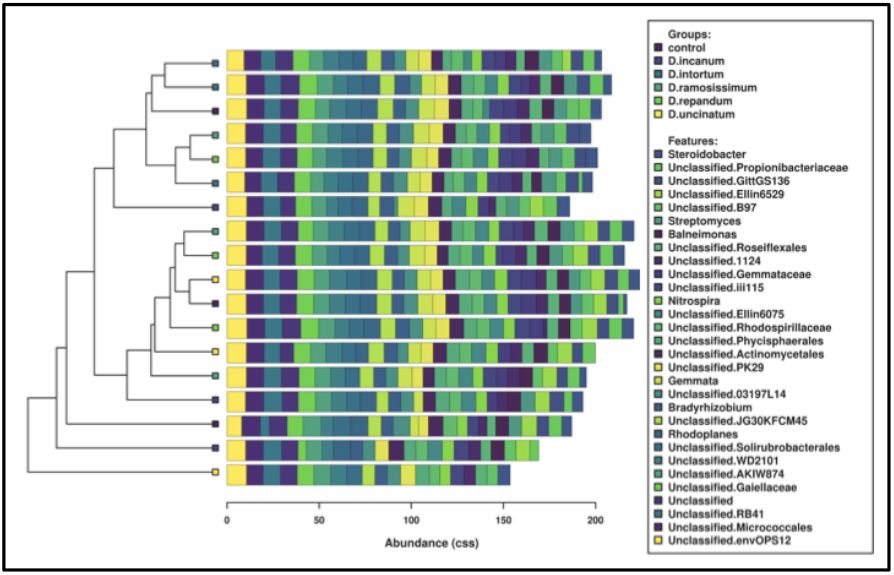
A clustered bar chart showing relative abundance of 30 most abundant soil bacterial genera in *Desmodium* spp. root zone soil and bulk soil. Most of the bacteria genera were unclassified and thus unidentified. The relative abundance was calculated from read counts (ASVs) normalized by total sum of squares (TSS) and transformed by cumulative sum scaling (CSS).

**Figure 3:**
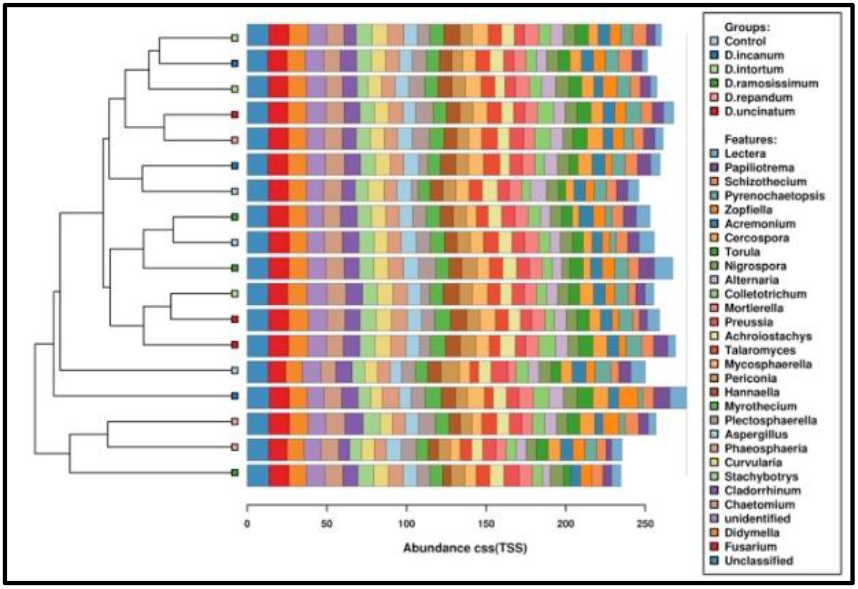
Clustered bar chart showing relative abundance of 30 most abundant soil fungal genera in *Desmodium* spp. root zone soil and bulk soil. Most of the fungal genera were classified and identified. The relative abundance was calculated from read counts (ASVs) normalized by total sum of squares (TSS) and transformed by cumulative sum scaling (CSS).

*Rhizobium, Bradyrhizobium* and *Mesorhizobium* (Xu et al., 2016). While investigating plant-endosymbionts relationships is important for plant health and productivity due to their intimate relationship with plant physiology, the importance of free living and the rhizosphere microbiome microorganisms cannot be overlooked, not the least because they are the source of the endophytes recruited by plants (Xiao et al., 2017). Free-living soil microbes also interact with plants through direct and indirect mechanisms that impact their health and productivity. In this study for example, some of the abundant bacterial groups identified are linked to varying activities in the soil that contribute to provision of ecosystem services. For example, *Nitrospira* spp. are known for their ability to fix nitrogen and potentially increasing supplies in the soil (Lu et al., 2020) while species of both *Streptomyces* and *Bradyrhizobium* are commonly known as biofertilizers (Htwe et al., 2019).

Contrary to expectations, fungal genera known to harbour plant pathogenic species were found in high abundance in both *Desmodium* plots and the bulk soil. These included *Fusarium, Gibberella* and *Didymella* genera (Figure 2). Although *Fusarium* is a ubiquitous genus with many harmless species, other species of this genus cause serious crop losses due to their pathogenicity and mycotoxin production that affect animals and human beings alike (Summerell, 2019). However, *Fusarium* species may form endophytic relationships with legumes such as *F. solani* and *Medicago truncatula* (Skiada et al., 2020) and become opportunistic when a favourable environment in the soil/plant is present.

In addition to *Fusarium*, we observed the presence of *Aspergillus* among the most abundant taxa in both *Desmodium* spp. plots and the bulk soil. Several species of the genus *Aspergillus* including *A. flavus, A. parasiticus* and *A. fumigatus* also produce potent mycotoxins that spoil cereal crop harvests and are harmful to human beings (Barkai-Golan, 2008). In our previous study, *Aspergillus* spp. were found in high abundance in soils of maize monoculture compared to long-term push-pull farms that employed *Desmodium* intercrops (Mwakilili et al., 2021). Similarly, (Maxwell et al., 2017) reported lower frequency of *Aspergillus flavus* in maize cobs from monoculture plots than in *Desmodium* intercropping push-pull systems, but an opposite trend for *A. parasiticus*. It is possible that the soils of the area are rich in these fungal taxa and the time under *Desmodium* spp. cultivation was too short to induce a significant change in populations abundance like in the discussed studies. In addition, without further analysis using higher resolution techniques such as whole genome sequencing (WGS) metagenomics, it is not possible to discern the specific *Fusarium* and *Aspergillus* species observed in the present study but the findings of this study point to a possibility of reduction of these taxa with continued cultivation of *Desmodium* spp.

The rest of the abundant genera were those ubiquitous in nature, containing beneficial, neutral and pathogenic fungi of plants and human beings, such as *Chaetomium, Cercophora, Colletotrichum* and *Plectosphaerella*.

### Common and unique soil microbial taxa among the *Desmodium* spp. plots

Comparison of the composition of soil microorganisms between the *Desmodium* spp. plots revealed the core microbiome of 29 bacteria and 55 fungi genera (Figure 4). Further, the microbiome and taxa that were uniquely associated with each *Desmodium* plot were identified. From the findings, the composition of taxa overlapped among the *Desmodium* plots, with *D. intortum* being associated with the largest number of unique bacterial genera (12) while *D. repandum* plots harboured the largest number of unique fungal genera (7). These two *Desmodium* spp. may be the most effective in recruiting and maintaining diverse microbial groups compared to others. Conversely, no unique bacterial genera were associated with the *D. incanum* plots (Figure 4A).

**Figure 4:**
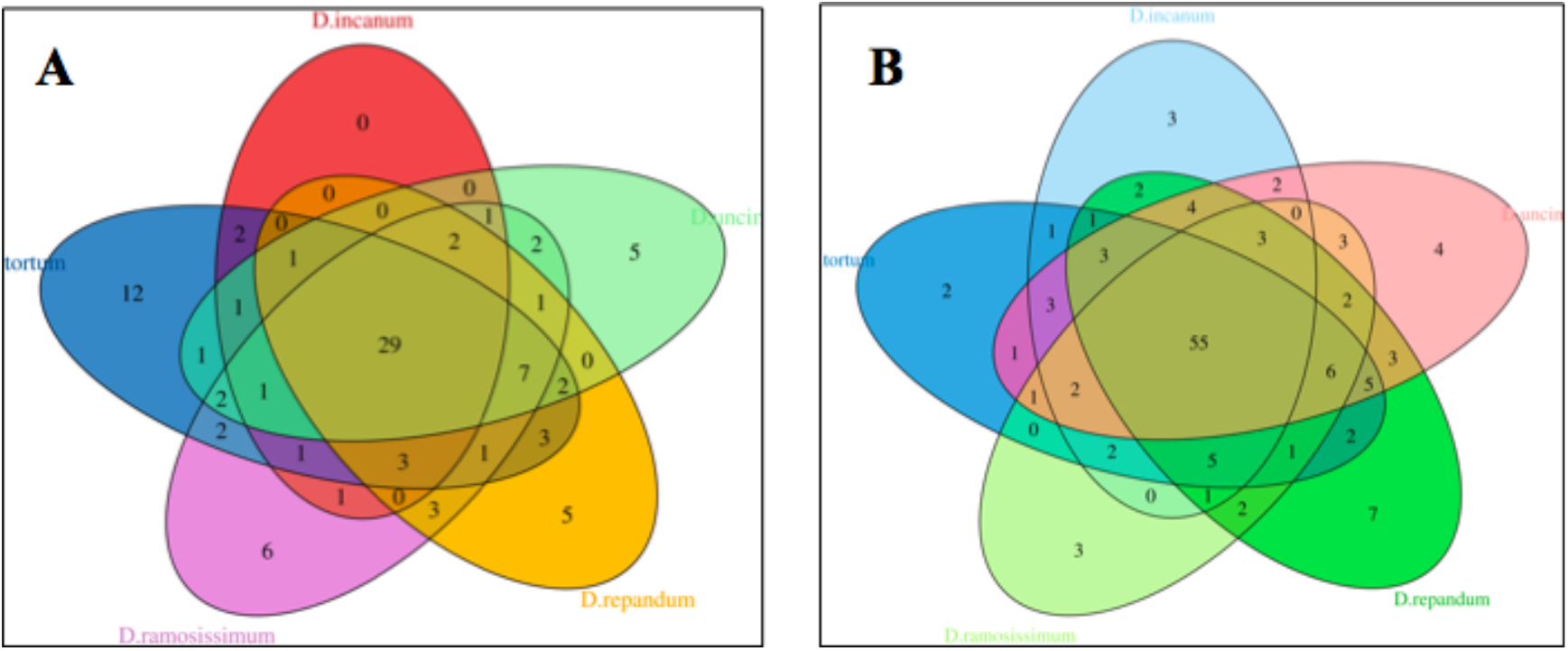
Venn diagram showing core, pan and unique soil bacterial (A) and fungal (B) genera from the 5 studied *Desmodium* species plots. The plots shared a large core genome of bacteria (29) and fungi (55) with few unique genomes associated with each *Desmodium*

Although the core fungal microbiome was larger than bacterial, most of the taxa were shared among the *Desmodium* plots causing the proportion of unique fungal taxa associated with individual *Desmodium* spp. plots to be lower. In general, most of the *Desmodium* spp. plots shared at least one taxon with each other, with *D. rammossisimum, D. repandum* and *D. intortum* sharing the largest number of both bacteria (7) and fungal genera (6) amongst themselves (Figure 4). This may indicate that their microbial recruitment strategies and root exudates composition are similar, possibly from a genetic makeup that is not very far from each other. The complete list of core, unique and pan genera is in supplementary tables 1 - 6.

While an association with more unique taxa in itself may not be an indication of direct and indirect activities of soil microbes that impact plant health, the ability of plants to recruit and support diverse microorganisms contributes to a more stable and resilient rhizospheric ecosystem (Wu et al., 2018). *D. intortum* and *D. uncinatum* are the commonly used intercrops in push-pull farming while the other *Desmodium* species have not been widely adopted despite some of them showing moderate to high drought tolerance, and *Striga* suppression. Of particular importance is *D. rammossisimum*, which along with *D. incanum*, showed the highest level of drought resistance and biomass retention in a previous field study (Midega et al., 2017). Although the study did not investigate the role of soil microbiome in the drought tolerance, other studies have demonstrated the ability of whole soil microbiomes to confer plants with the ability to tolerate abiotic stresses including tolerance to drought (Zolla et al., 2013; Vurukonda et al., 2016; Huang et al., 2017). In addition to finding the link between belowground diversity and abiotic stress tolerance, it may be useful to investigate the potential of mixed-intercropping in push-pull systems by combining more than one *Desmodium* spp. to leverage both below- and above-ground benefits offered by different species.

### Diversity measures

We also analysed different measures of diversity and richness of the soil microbiome among the *Desmodium* spp. plots and the bulk soil. Comparing the diversity of soil microbes within each treatment (alpha diversity), we found no significant difference/variation of both bacterial (Supplementary figure 1) and fungal (Supplementary figure 2) communities through diversity measures of richness and evenness.

Similarly, analysis of diversity between the treatments (beta diversity) did not reveal any significant association of the soil microbial populations to the different treatments i.e. *Desmodium* spp. plots or the bulk soil. This indicates the overall variation of the soil microbial communities composition between plots was random and not significantly altered by the cultivation of *Desmodium* spp. compared to the bulk soil, as observed by absence of distinct clustering patterns in PCoA plots (Figure 5).

**Figure 5:**
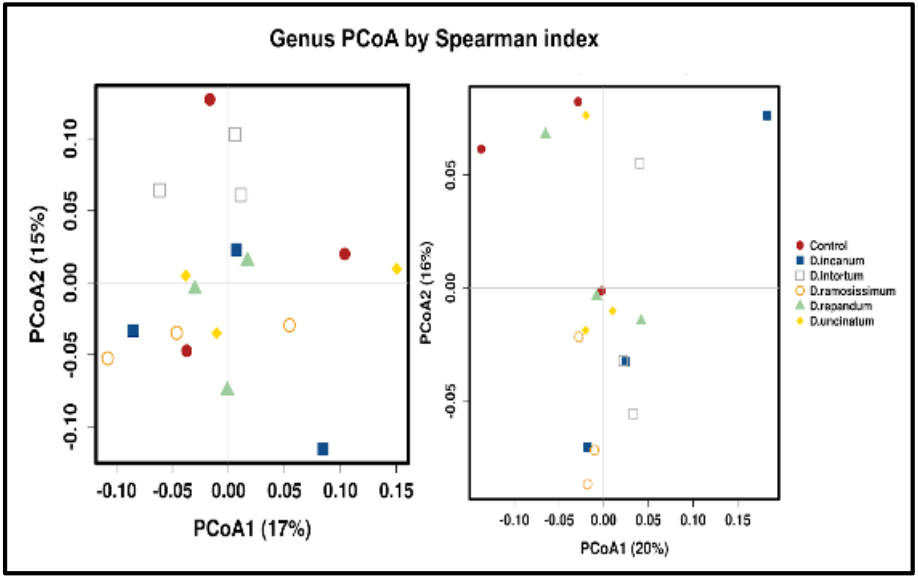
PCoA plots of soil bacterial (left) and fungal (right) genera (OTUs) across the *Desmodium* spp. root zone soil and bulk soil based on Spearman correlation. The clustering patterns reveal no association between the different treatments and microbial populations, indicating no clear impact of the *Desmodium* species.

We expected to see a more pronounced impact of the *Desmodium* spp. cultivation on the diversity and divergence of soil microbial communities compared to bulk soil. Our observations suggest that two years is not a sufficient time window for a noticeable influence of *Desmodium* spp. on whole shifts in belowground microbial communities. In a previous study, we reported the impact of long term (14 - 18 years) *Desmodium* intercropping on the composition and diversity of soil microbial profiles (Mwakilili et al., 2021) where a strong shift of fungal communities was observed in push-pull plots compared to maize monoculture plots.

Other studies have indeed suggested that microbial-based plant-soil feedback is a slow process in that although aboveground vegetation has the largest influence on assemblages and alterations of soil microbial communities, the process may take several years to form stable structures (Eisenhauer et al., 2011; Vukicevich et al., 2016). Given a longer period, the patterns of the impact of the *Desmodium* spp. on soil microbial communities may emerge, and with them, other emergent differential benefits conferred by soil microorganisms on *Desmodium* plant health and other ecological services.

### Differential abundances of individual taxa

Despite the lack of significant difference in overall diversity and richness of the soil microbial populations, several bacterial and fungal taxa were enriched in specific *Desmodium* spp. plots as well as in the bulk soil (Figure 6 and 7). A larger proportion of the significantly abundant taxa are fungal (23) compared to only four bacterial taxa. Of the four significantly abundant bacteria taxa, one taxon code-named JG30KFCM45 was abundant in all treatments. By contrast, the genus *Agromyces* was significantly abundant in *D. intortum* plots only (Figure 7). *Novosphingobium* and *Craurococcus* were other significantly abundant bacterial genera, both having high abundance in *D. repandum* plots. *Novosphingobium* was in addition found in significantly higher abundance in *D. intortum* plots and *Craurococcus* in *D. uncinatum* plots and the bulk soil.

**Figure 6:**
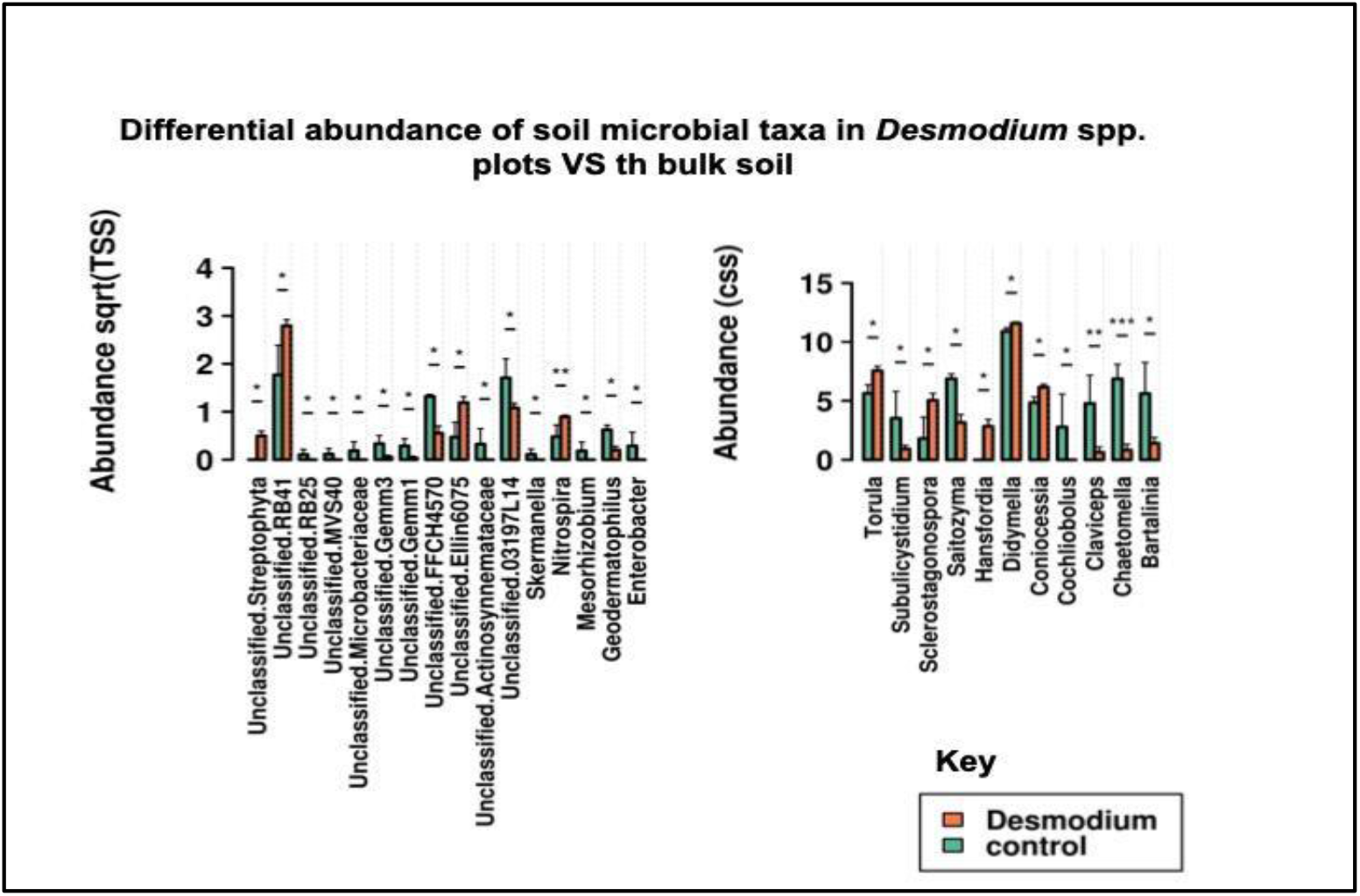
Comparison of differential abundance of bacterial (left) and fungal (right) taxa between *Desmodium* spp. root zone soil and bulk soil highlighting taxa significantly abundant in either treatment. (ANOVA, where * = p<0.05, ** = p<0.01, *** = p<0.001)

**Figure 7:**
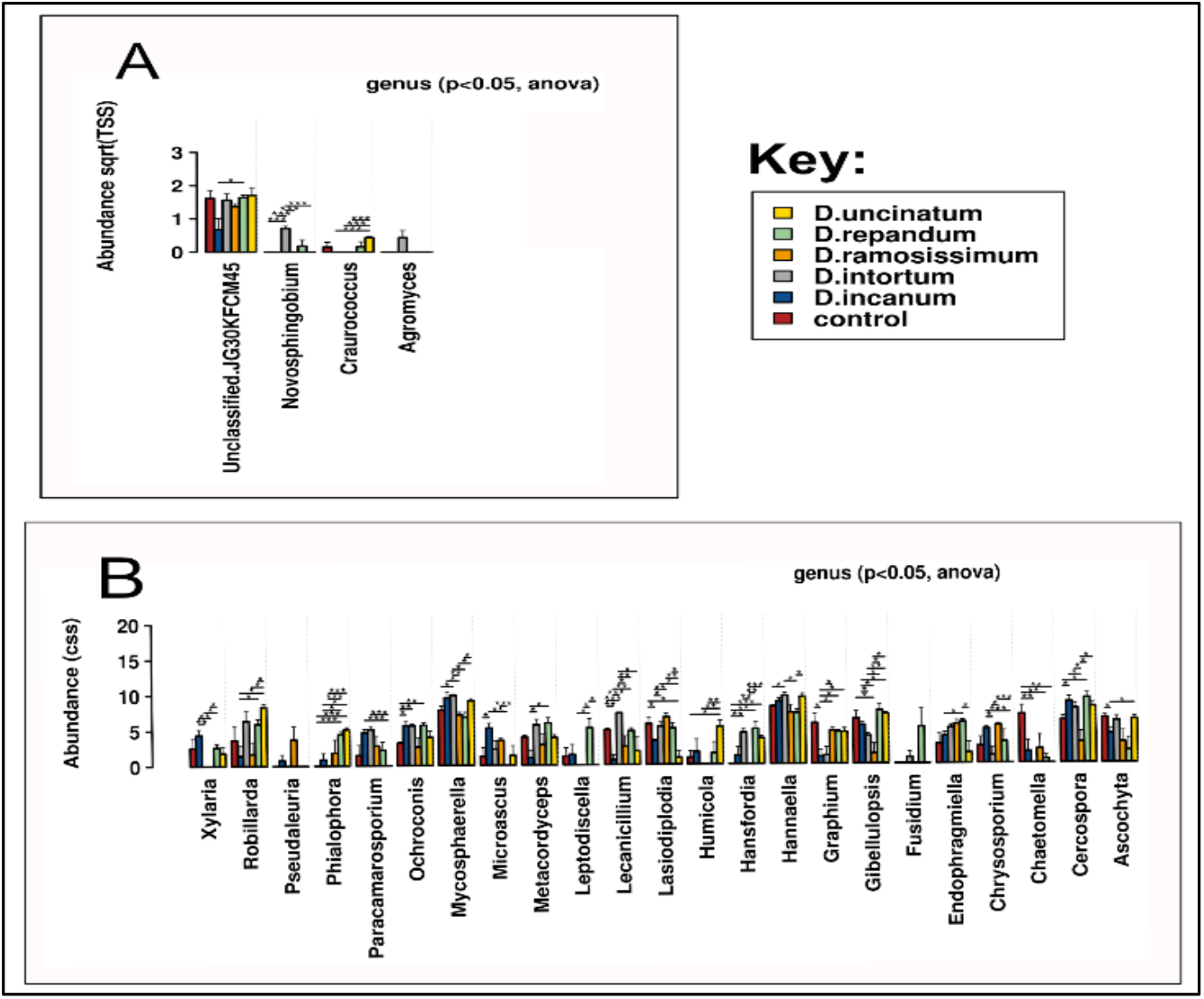
A bar chart showing significantly abundant bacterial (A) and fungal (B) taxa among *Desmodium* spp. root zone soil and bulk soil obtained by t-test pairwise comparisons (where *p<0.05, **p<0.01, ***p<0.001). Error bars depict standard error.

Among the fungal taxa that were significantly abundant, 12 of them were the most common being significantly abundant across most of the treatments. *Mycosphaerella, Hannaella* and *Cercospora* were the most ubiquitous significantly abundant fungal taxa irrespective of treatment. On the other hand, other fungal groups were significantly abundant in few treatments such as *Pseudaleuria* and *Fusidum*, which were enriched in only two treatments (*D. incanum* and *D. ramosissimum)*. In addition, three of the five *Desmodium* spp. plots harboured a substantial percentage of the significantly abundant taxa; these are *D. uncinatum, D. incanum* and *D. intortum* (Figure 7.)

As in our previous long-term study on maize-*Desmodium* intercropping soils (Mwakilili et al., 2021) where we investigated impact of push-pull technology that employs perennial *Desmodium* spp. intercrops, soils diverged more in fungal than bacterial communities. The studied plots were between 14 and 18 years old and employed *D. intortum* and/or *D. uncinatum* as perennial intercrops. To our knowledge, there are no other studies that investigate the impact of *Desmodium* cropping on whole soil microbial populations. Understanding which shifts in soil microbiomes are associated with plant health and productivity is, however, important to be able to more efficiently reap the benefits of ecologically intensified cropping systems.

Without experimental verification, we can only speculate about the role of and potential ecosystem services rendered by differentially abundant taxa in *Desmodium* plots. The bacterial genera *Novosphingobium* and *Agromyces*, both significantly enriched in *Desmodium* spp. plots, harbour species with the ability to degrade complex organic compounds, with the latter also being able to resist heavy metals. Members of the taxa have been reported to show capability of degrading complex organic compounds such as xylan (Rivas et al., 2004) and aromatic compounds (Sohn et al., 2004; Liu et al., 2005). These properties may be essential in releasing nutrients from complex organic matter as well as degrading toxic compounds and thus potentially improve the ability of plants to survive in harsh environments.

Several fungal taxa were significantly abundant in *Desmodium* plots but not in the bulk soil, including *Pseudaleuria, Phialophora, Hansfordia* and *Fusidium*. These genera comprise common soil and wood saprotrophs most of which have known ecological functions. The genus *Pseudaleuria* for instance has been associated with healthy soils and disease suppression in pea fields (Xu et al., 2012), while some species of *Phialophora* cause soft rot of wood and other root diseases especially in wheat and other species show plant protection properties (Zriba et al., 1999; Karunasekera & Daniel, 2013). The remaining fungal genera that were significantly abundant in *Desmodium* spp. plots belong to endophytic groups, and often possess beneficial plant protection and growth promotion activity. These include *Hannaella*, a genus of endophytic fungi (Gonzaga et al., 2015), *Chrysosporium*, whose species have been shown to produce plant hormones (Hamayun et al., 2009), and *Lecanicillium*, a genus comprising species that display a wide range of growth promotion and protection activities against pathogens, insects and nematodes on plants (Goettel et al., 2008; Nicoletti & Becchimanzi, 2020). Other fungal groups highly abundant in *Desmodium* plots, but present in the bulk soil in small amounts were *Paracamarosporium, Microasus, Leptodiscella* and *Humicola*.

Conversely, species of some fungal genera that were significantly abundant in the bulk soil compared to the *Desmodium* plots (i.e. *Ascochyta, Chaetomella* and *Graphium*) have been reported to cause diseases in plants. For example, species of the genus *Ascochyta*, a teleomorph of *Didymella* spp., cause blights of cereals and legumes (Tivoli & Banniza, 2007)

Although not conclusive, these findings point in a slow divergence, whereby cultivation of *Desmodium* spp. favours growth and replication of specific groups of microbial taxa. Most of the microbial groups found in significantly higher abundance in *Desmodium* spp. plots are either ubiquitous harmless microbes or have previously been noted for conferring ecosystem services such as improved access to nutrients from the soil, suppressing harmful and disease-causing microbes. Indeed, reports of lower mycotoxin producing fungi in push-pull plots where *Desmodium* spp. are used as intercrops (Maxwell et al., 2017; Owuor et al., 2018; Njeru et al., 2020) as well as associational resistance in maize grown on soil from long term push-pull fields (Mutyambai et al., 2019) may be the first clue about the important role of *Desmodium* spp. in shaping soil microbial communities leading to diverse ecological benefits related to food production and safety. These observations warrant further dissection.

## CONCLUSION

In this study, we hypothesised that continuous cultivation of *Desmodium* species shifts in soil microbial populations structure, composition and diversity relative to non-cultivated bulk soil. In addition, we hypothesised that the impact on soil microbial communities would be dif ferent among different *Desmodium* species. Although cultivation of *Desmodium* spp. leads to significant increases in abundance of selected bacterial and fungal taxa, no significant difference in overall diversity of soil microbial communities both within plots and between plots. Soil microbial communities interact with plants and play a key role in restoring resilience of soils for provision of ecosystem services in farming systems.

However, as shown in this study, shifts in microbial populations are more intricate long-term processes than anticipated without short-term incentives. A longer period of cultivation is undoubtedly required for clearer patterns of changes in the composition and abundance of the soil microbial communities. Aboveground vegetation has been demonstrated to play the most significant role in shaping soil microbial communities in long-term studies. This fits well with the nature of push-pull farming, being a perennial *Desmodium* spp. based intercropping technology whose numerous benefits become apparent with time, adding to unseen belowground ecological services of the technology.

## ACKNOWLEDGEMENT

We are grateful for the support of ICIPE staff during soil sample collection and processing, including Dickens Nyagol, Levi and Moureen. Special thanks to Emma Huri, Will English and Paul Eagan for reviewing early drafts of the manuscript.

## Supplementary materials

### Supplementary figures

**Supplementary figure 1:**
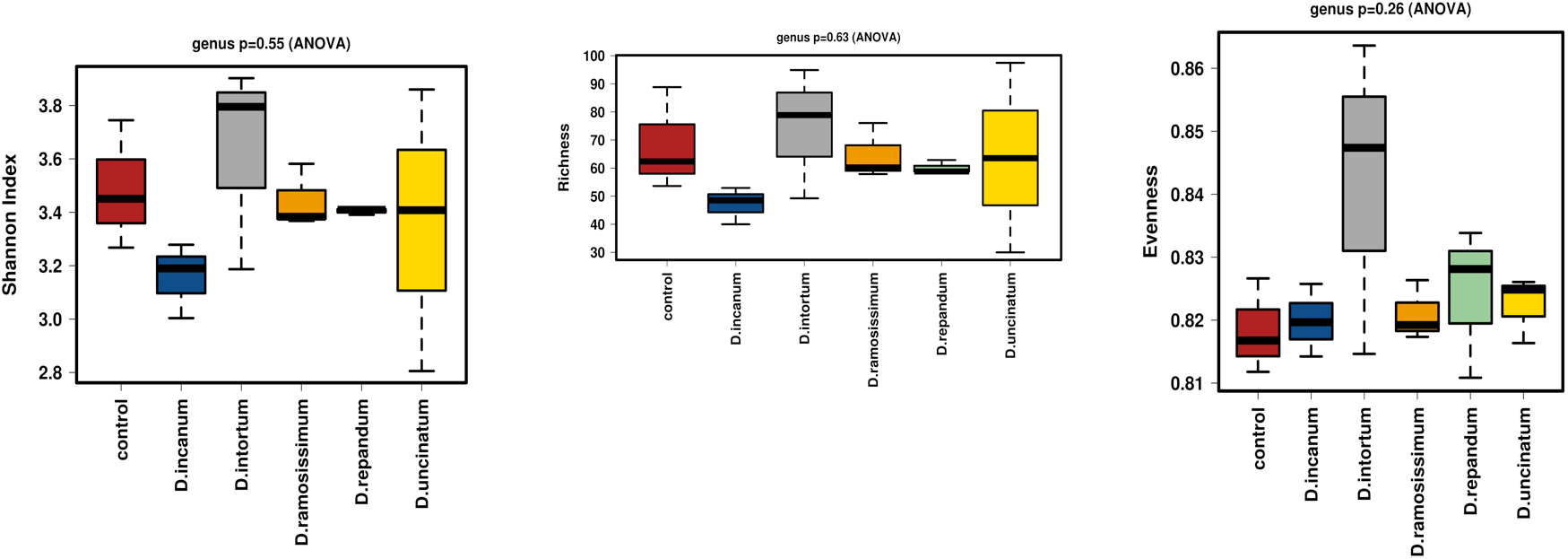
Box plot representation of alpha diversity measures (from left, Shannon index, richness and evenness measures) of soil bacterial communities in all treatments. Alpha diversity measures compare diversity of microbial populations within each treatment. The figures show variation in the diversity in composition of soil bacterial taxa (ANOVA, p = 0.55) in the treatments was not significant. In addition, there was no significant difference in richness (ANOVA, p = 0.63) and evenness of the taxa (ANOVA, p = 0.26).

**Supplementary figure 2:**
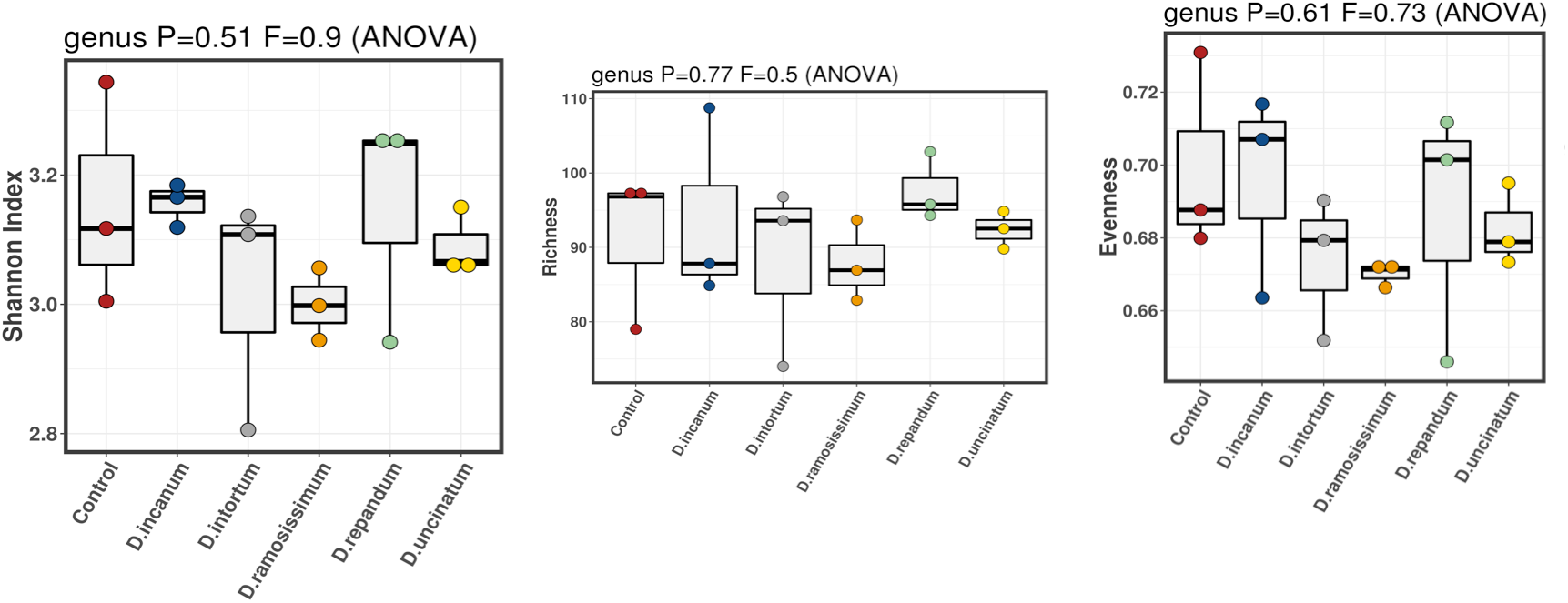
Box plot representation of alpha diversity measures (from left, Shannon index, richness and evenness measures) of soil fungal taxa in all treatments. Alpha diversity measures compare diversity of microbial populations within each treatment. The figures show variation in the diversity in composition of soil fungal taxa (ANOVA, p = 0.51) in the treatments was not significant. The same observation was made for richness (ANOVA, p = 0.77) and evenness (ANOVA, p = 0.61) measures.

### Supplementary tables

**Supplementary table 1:** Unique bacterial taxa that occur in only one of the *Desmodium* spp. plots

| Taxa | Abundance |  |  |  |  | OCC |  |  |  |  |
| --- | --- | --- | --- | --- | --- | --- | --- | --- | --- | --- |
|  | *D. incanum* | *D. intortum* | *D. ramosissimum* | *D. repadum* | *D. uncinatum* | *D. incanum* | *D. intortum* | *D. ramosissimum* | *D. repadum* | *D. uncinatum* |
| ***D. repadum*** |  |  |  |  |  |  |  |  |  |  |
| *X5B12* | 0 | 0 | 1.697 | 3.267 | 2.043 | 0 | 0 | 0.33 | 0.67 | 0.33 |
| *X03196A21* | 0 | 0 | 0.887 | 2.137 | 0.373 | 0 | 0 | 0.33 | 0.67 | 0.33 |
| *Unclassified CV90* | 1.62 | 0 | 0 | 2.483 | 2.02 | 0.33 | 0 | 0 | 0.67 | 0.33 |
| *OM27* | 1.79 | 1.69 | 0 | 2.347 | 1.227 | 0.33 | 0.33 | 0 | 0.67 | 0.33 |
| *Dolo 23* | 0 | 1.837 | 2.013 | 4.357 | 2.07 | 0 | 0.33 | 0.33 | 0.67 | 0.33 |
| ***D. intortum*** |  |  |  |  |  |  |  |  |  |  |
| *Unclassified TM73* | 1.363 | 3.15 | 0 | 0 | 0 | 0.33 | 0.67 | 0 | 0 | 0 |
| *Unclassified TM71* | 0 | 3.087 | 0 | 1.533 | 1.49 | 0 | 0.67 | 0 | 0.33 | 0.33 |
| *Unclassified TM7* | 0 | 2.867 | 1.697 | 0 | 0.84 | 0 | 0.67 | 0.33 | 0 | 0.33 |
| *Unclassified B07_WMSP1* | 0 | 3.39 | 1.057 | 2.193 | 2.127 | 0 | 0.67 | 0.33 | 0.33 | 0.33 |
| *Unclassified Acidimicrobiales* | 0 | 4.227 | 0 | 0 | 0 | 0 | 0.67 | 0 | 0 | 0 |
| *Sphingomonadaceae* | 0 | 6.11 | 0 | 1.533 | 2.257 | 0 | 1 | 0 | 0.33 | 0.33 |
| *Rhizobiaceae* | 2.027 | 6.79 | 2.007 | 2.143 | 1.49 | 0.33 | 1 | 0.33 | 0.33 | 0.33 |
| *Pseudomonadaceae* | 0 | 2.993 | 0 | 0 | 1.147 | 0 | 0.67 | 0 | 0 | 0.33 |
| *Micrococcaceae* | 0 | 3.843 | 1.877 | 0 | 0 | 0 | 0.67 | 0.33 | 0 | 0 |
| *Microbacteriaceae* | 0 | 4.287 | 1.74 | 1.727 | 1.467 | 0 | 0.67 | 0.33 | 0.33 | 0.33 |
| *Chloroflexaceae* | 0 | 2.663 | 0 | 0 | 0 | 0 | 0.67 | 0 | 0 | 0 |
| *Entothionellaceae* | 1.53 | 1.237 | 0.977 | 0.727 | 0 | 0.33 | 0.67 | 0.33 | 0.33 | 0 |
| ***D. uncinatum*** |  |  |  |  |  |  |  |  |  |  |
| *Unclassified S085* | 2.313 | 0 | 1.597 | 1.927 | 3.147 | 0.33 | 0 | 0.33 | 0.33 | 0.67 |
| *Unclassified C0119* | 0 | 1.317 | 1.27 | 0.873 | 2.5 | 0 | 0.33 | 0.33 | 0.33 | 0.67 |
| *Acetobacteraceae* | 0 | 0 | 0 | 1.623 | 4.167 | 0 | 0 | 0 | 0.33 | 1 |
| *Haliangiaceae* | 1.217 | 0 | 0 | 0 | 2.893 | 0.33 | 0 | 0 | 0 | 0.67 |
| *Flavobacteriaceae* | 1.91 | 0 | 0 | 0.93 | 3 | 0.33 | 0 | 0 | 0.33 | 0.67 |
| ***D. ramosissimum*** |  |  |  |  |  |  |  |  |  |  |
| *Unclassified Planctomycetes* | 0 | 1.767 | 3.38 | 1.227 | 2.03 | 0 | 0.33 | 0.67 | 0.33 | 0.33 |
| *Geodermatophilaceae* | 1.97 | 0 | 3.203 | 1.58 | 1.467 | 0.33 | 0 | 0.67 | 0.33 | 0.33 |
| *Unclassified CL50015* | 0 | 0 | 2.707 | 0 | 1.863 | 0 | 0 | 0.67 | 0 | 0.33 |
| *Unclassified CCU21* | 0 | 1.64 | 2.953 | 0 | 0 | 0 | 0.33 | 0.67 | 0 | 0 |
| *Ardenscatenaceae* | 0 | 0 | 2.397 | 0 | 0 | 0 | 0 | 0.67 | 0 | 0 |
| *Cystobacteraceae* | 0 | 1.84 | 3.403 | 1.667 | 1.993 | 0 | 0.33 | 0.67 | 0.33 | 0.33 |

**Supplementary table 2:** Core bacterial taxa found in all five *Desmodium* spp. Plots

| Taxa | Abundance |  |  |  |  | OCC |  |  |  |  |
| --- | --- | --- | --- | --- | --- | --- | --- | --- | --- | --- |
|  | *D. incanum* | *D. intortum* | *D. ramosissimum* | *D. repandum* | *D. uncinatum* | *D. incanum* | *D. intortum* | *D. ramosissimum* | *D. repandum* | *D. uncinatum* |
| *Unclassified WD2101* | 6.907 | 8.047 | 8.003 | 7.903 | 9.037 | 1 | 1 | 1 | 1 | 1 |
| *Unclassified TK10* | 3.433 | 6.087 | 5.48 | 6.197 | 3.84 | 0.67 | 1 | 1 | 1 | 0.67 |
| *Unclassified Solirubrobacterales* | 4.977 | 8.123 | 8.12 | 8.103 | 7.993 | 0.67 | 1 | 1 | 1 | 1 |
| *Unclassified Roseiflexales* | 3.82 | 5.517 | 4.273 | 5.79 | 6.367 | 0.67 | 1 | 0.67 | 1 | 1 |
| *Unclassified RB41* | 9.843 | 8.983 | 9.593 | 9.55 | 9.147 | 1 | 1 | 1 | 1 | 1 |
| *Unclassified PK29* | 5.047 | 7.01 | 7.073 | 6.903 | 7.213 | 0.67 | 1 | 1 | 1 | 1 |
| *Unclassified Phycisphaerales* | 6.607 | 5.78 | 6.207 | 6.507 | 7.523 | 1 | 1 | 1 | 1 | 1 |
| *Unclassified Myxococcales* | 3.367 | 5.403 | 5.347 | 4.84 | 4.157 | 0.67 | 1 | 1 | 0.67 | 0.67 |
| *Unclassified Micrococcales* | 9.463 | 9.267 | 9.503 | 9.17 | 9.493 | 1 | 1 | 1 | 1 | 1 |
| *Unclassified JG30KFCM45* | 4.28 | 7.8 | 7.397 | 8.03 | 7.997 | 0.67 | 1 | 1 | 1 | 1 |
| *Unclassified.envOPS12* | 10.257 | 9.947 | 10.337 | 10.513 | 10.44 | 1 | 1 | 1 | 1 | 1 |
| *Unclassified.B97* | 4.1 | 3.87 | 5.86 | 6.573 | 5.81 | 0.67 | 0.67 | 1 | 1 | 1 |
| *Unclassified Actinomycetales* | 7.403 | 6.567 | 6.5 | 6.847 | 4.71 | 1 | 1 | 1 | 1 | 0.67 |
| *Unclassified I124* | 5.327 | 5.573 | 6.08 | 4.877 | 5.98 | 1 | 1 | 1 | 1 | 1 |
| *Unclassified 03197L14* | 5.397 | 6.95 | 6.647 | 6.85 | 6.8 | 1 | 1 | 1 | 1 | 1 |
| *Unclassified* | 7.76 | 8.997 | 8.553 | 8.907 | 8.367 | 1 | 1 | 1 | 1 | 1 |
| *Sinobacteraceae* | 5.673 | 4.927 | 5.863 | 6.597 | 5.897 | 1 | 1 | 1 | 1 | 1 |
| *Rhodospirillaceae* | 4.283 | 6.443 | 6.507 | 6.777 | 6.343 | 0.67 | 1 | 1 | 1 | 1 |
| *Propionibacteriaceae* | 7.827 | 7.4 | 7.213 | 7.32 | 7.303 | 1 | 1 | 1 | 1 | 1 |
| *Pirellulaceae* | 5.007 | 2.693 | 7.127 | 3.753 | 3.837 | 0.67 | 0.67 | 1 | 0.67 | 0.67 |
| *Nitrospiraceae* | 6.353 | 6.003 | 5.727 | 6.257 | 6.477 | 1 | 1 | 1 | 1 | 1 |
| *mb2424* | 4.46 | 4.063 | 4.283 | 3.973 | 3.92 | 0.67 | 0.67 | 0.67 | 0.67 | 0.67 |
| *Hyphomicrobiaceae* | 9 | 8.623 | 5.86 | 8.547 | 5.53 | 1 | 1 | 0.67 | 1 | 0.67 |
| *Gemmataceae* | 5.153 | 7.9 | 8.437 | 8.873 | 5.623 | 0.67 | 1 | 1 | 1 | 0.67 |
| *Gaiellaceae* | 7.563 | 9.32 | 8.613 | 9.243 | 9.26 | 1 | 1 | 1 | 1 | 1 |
| *Ellin6075* | 7.76 | 6.017 | 7.217 | 7.143 | 5.01 | 1 | 1 | 1 | 1 | 0.67 |
| *Chitinophagaceae* | 7.147 | 6.767 | 7 | 6.837 | 4.51 | 1 | 1 | 1 | 1 | 0.67 |
| *Bradyrhizobiaceae* | 7.75 | 8.513 | 7.817 | 7.77 | 7.787 | 1 | 1 | 1 | 1 | 1 |
| *AKIW874* | 8.443 | 8.107 | 8.443 | 8.217 | 8.647 | 1 | 1 | 1 | 1 | 1 |

**Supplementary table 3:** Pan bacterial taxa shared by several *Desmodium* spp. plots

| Taxa | *Desmodium* spp.<br>plots sharing taxa | Abundance |  |  |  |  | OCC |  |  |  |  |
| --- | --- | --- | --- | --- | --- | --- | --- | --- | --- | --- | --- |
|  |  | *D.*<br>*incanum* | *D.*<br>*intortum* | *D.*<br>*ramosissimum* | *D.*<br>*repandum* | *D.*<br>*uncinatum* | *D.*<br>*incanum* | *D.*<br>*intortum* | *D.*<br>*ramosissimum* | *D.*<br>*repandum* | *D.*<br>*uncinatum* |
| Unclassified *Streptophyta* | DIN, DIT, DRP, DUN | 4.437 | 4.617 | 1.717 | 3.997 | 4.38 | 0.67 | 1 | 0.33 | 0.67 | 1 |
| Unclassified *Sphingobacteriales* | DIN, DIT | 3.05 | 1.727 | 1.17 | 0 | 0 | 0.67 | 0.67 | 0.33 | 0 | 0 |
| Unclassified S0208 | DIT, DRM, DUN | 0 | 3.14 | 3.497 | 1.917 | 3.5 | 0 | 0.67 | 0.67 | 0.33 | 0.67 |
| Unclassified *Pla4* | DRM, DRP | 0 | 1.257 | 4.253 | 5.897 | 1.96 | 0 | 0.33 | 0.67 | 1 | 0.33 |
| Unclassified MVP88 | DIN, DIT, DRM | 2.75 | 2.457 | 2.937 | 1.597 | 1.407 | 0.67 | 0.67 | 0.67 | 0.33 | 0.33 |
| Unclassified MND1 | DIN, DRM | 4.293 | 1.687 | 4.137 | 2.47 | 1.863 | 0.67 | 0.33 | 0.67 | 0.33 | 0.33 |
| Unclassified iii115 | DIT, DRM, DRP, DUN | 2.677 | 6.84 | 6.697 | 6.417 | 5.007 | 0.33 | 1 | 1 | 1 | 0.67 |
| Unclassified H39 | DRM, DRP | 1.75 | 1.393 | 3.03 | 2.95 | 1.623 | 0.33 | 0.33 | 0.67 | 0.67 | 0.33 |
| Unclassified GittGS136 | DIT, DRM, DRP, DUN | 2.55 | 6.823 | 6.343 | 6.283 | 4.24 | 0.33 | 1 | 1 | 1 | 0.67 |
| Unclassified *Gemmatimonadetes* | DRM, DUN | 0 | 1.213 | 1.82 | 0 | 3.057 | 0 | 0.33 | 0.67 | 0 | 0.67 |
| Unclassified Ellin6529 | DIN, DRM, DRP, DUN | 7.59 | 1.66 | 5.15 | 5.513 | 5.263 | 1 | 0.33 | 0.67 | 0.67 | 0.67 |
| Unclassified Ellin329 | DRM, DUN | 0 | 0 | 3.357 | 1.767 | 2.71 | 0 | 0 | 0.67 | 0.33 | 0.67 |
| Unclassified DRC31 | DIN, DIT, DRM, DUN | 3.667 | 4.293 | 3.73 | 2.157 | 4.273 | 0.67 | 0.67 | 0.67 | 0.33 | 0.67 |
| Unclassified *Bacteria* | DIN, DRM, DRP, DUN | 3.303 | 2.193 | 4.293 | 4.367 | 3.067 | 0.67 | 0.33 | 1 | 1 | 0.67 |
| Unclassified AKIW781 | DIT, DRM | 1.583 | 4.997 | 4.923 | 1.533 | 1.883 | 0.33 | 0.67 | 0.67 | 0.33 | 0.33 |
| Unclassified agg27 | DRM, DRP | 1.583 | 0 | 4.717 | 2.453 | 1.85 | 0.33 | 0 | 1 | 0.67 | 0.33 |
| Unclassified *Acidobacteria5* | DIN, DRM, DUN | 3.303 | 1.513 | 3.147 | 1.73 | 3.843 | 0.67 | 0.33 | 0.67 | 0.33 | 0.67 |
| Unclassified ABY1 | DIN, DIT, DRM, DRP | 3.167 | 3.03 | 4.493 | 2.507 | 1.787 | 0.67 | 0.67 | 1 | 0.67 | 0.33 |
| Unclassified 03196E2 | DIT, DRM, DRP, DUN | 0 | 4.723 | 3.32 | 2.847 | 3.363 | 0 | 1 | 0.67 | 0.67 | 0.67 |
| *Streptomycetaceae* | DIN, DIT, DRM, DRP | 4.877 | 7.603 | 4.927 | 4.86 | 2.07 | 0.67 | 1 | 0.67 | 0.67 | 0.33 |
| *Sporichthyaceae* | DIT, DUN | 2.383 | 6 | 1.91 | 2.12 | 3.857 | 0.33 | 1 | 0.33 | 0.33 | 0.67 |
| *Solirubrobacteraceae* | DIT, DRM, DRP | 1.513 | 4.36 | 3.967 | 4.803 | 2.327 | 0.33 | 0.67 | 0.67 | 0.67 | 0.33 |
| RB40 | DRM, DRP, DUN | 0 | 0 | 2.5 | 3.013 | 2.303 | 0 | 0 | 0.67 | 0.67 | 0.67 |
| *Pseudonocardiaceae* | DIT, DRM, DRP, DUN | 2.077 | 6.127 | 5.44 | 3.947 | 4.13 | 0.33 | 1 | 1 | 0.67 | 0.67 |
| *Planctomycetaceae* | DIT, DRP, DUN | 1.74 | 5.013 | 1.727 | 5.023 | 3.34 | 0.33 | 1 | 0.33 | 1 | 0.67 |
| *Oxalobacteraceae* | DIT, DRP | 0 | 5.167 | 1.407 | 3.857 | 0 | 0 | 1 | 0.33 | 1 | 0 |
| *Nocardiodaceae* | DIT, DRM, DUN | 1.237 | 6.503 | 3.963 | 1.417 | 3.803 | 0.33 | 1 | 0.67 | 0.33 | 0.67 |
| mitochondria | DIT, DRP | 2.067 | 3.203 | 1.277 | 3.69 | 1.267 | 0.33 | 0.67 | 0.33 | 0.67 | 0.33 |
| *Micromonosporaceae* | DIN, DIT, DRM, DRP | 4.697 | 5.693 | 5.28 | 5.023 | 2.49 | 0.67 | 0.67 | 0.67 | 0.67 | 0.33 |
| *Kouleothrixaceae* | DIT, DRM, DRP, DUN | 2.14 | 3.95 | 3.537 | 4.087 | 6.123 | 0.33 | 0.67 | 0.67 | 0.67 | 1 |
| FCH4570 | DIT, DRP, DUN | 2.21 | 4.333 | 2.17 | 3.897 | 4.82 | 0.33 | 0.67 | 0.33 | 0.67 | 0.67 |
| *Euzebyaceae* | DIN, DIT | 3.69 | 2.583 | 1.347 | 0 | 1.467 | 0.67 | 0.67 | 0.33 | 0 | 0.33 |
| *Cytophagaceae* | DIN, DIT, DUN | 3.51 | 3.41 | 1.74 | 0 | 3.723 | 0.67 | 0.67 | 0.33 | 0 | 0.67 |
| *Comamonadaceae* | DIT, DRM, DRP, DUN | 0 | 3.76 | 3.087 | 3.107 | 3.11 | 0 | 0.67 | 0.67 | 0.67 | 0.67 |
| *Caldilineaceae* | DIT, DRM | 0 | 3.097 | 2.77 | 1.727 | 1.693 | 0 | 0.67 | 0.67 | 0.33 | 0.33 |
| *Bacillaceae* | DIT, DRP | 1.91 | 2.853 | 1.757 | 3.183 | 1.66 | 0.33 | 0.67 | 0.33 | 0.67 | 0.33 |
| *A4b* | DIT, DRM, DRP, DUN | 1.91 | 6.167 | 3.937 | 2.95 | 6.837 | 0.33 | 1 | 0.67 | 0.67 | 1 |

**Legend:** DRM = *D. ramosissimum*, DUN = *D. uncinatum*, DIN = *D. incanum*, DIT = *D. intortum*, DRP = *D. repandum*

**Supplementary table 4:** Unique soil fungal genera in *Desmodium* spp. plots

|  | Abundance |  |  |  |  | OCC |  |  |  |  |
| --- | --- | --- | --- | --- | --- | --- | --- | --- | --- | --- |
| Taxa | *D. incanum* | *D. intortum* | *D. ramosissimum* | *D. repandum* | *D. uncinatum* | *D. incanum* | *D. intortum* | *D. ramosissimum* | *D. repandum* | *D. uncinatum* |
| ***D. intortum*** |  |  |  |  |  |  |  |  |  |  |
| *Zygosporium* | 0 | 2.007 | 0 | 2.213 | 0 | 0 | 0.67 | 0 | 0.33 | 0 |
| *Knufia* | 2.563 | 2.907 | 1.35 | 0 | 0 | 0.33 | 0.67 | 0.33 | 0 | 0 |
| ***D. ramosissimum*** |  |  |  |  |  |  |  |  |  |  |
| *Stephanonectria* | 1.677 | 0 | 3.767 | 1.25 | 0 | 0.33 | 0 | 0.67 | 0.33 | 0 |
| *Solheimia* | 1.377 | 0 | 2.31 | 0 | 0 | 0.33 | 0 | 0.67 | 0 | 0 |
| *Pseudaleuria* | 0.787 | 0 | 3.71 | 0 | 0 | 0.33 | 0 | 0.67 | 0 | 0 |
| ***D. repandum*** |  |  |  |  |  |  |  |  |  |  |
| *Fusidium* | 0 | 0.823 | 0 | 5.15 | 0 | 0 | 0.33 | 0 | 0.67 | 0 |
| *Atractiella* | 1.887 | 0 | 0 | 3.71 | 0 | 0.33 | 0 | 0 | 0.67 | 0 |
| *Veronaea* | 0.68 | 0 | 0 | 3.207 | 0 | 0.33 | 0 | 0 | 0.67 | 0 |
| *Veronaea* | 0.68 | 0 | 0 | 3.207 | 0 | 0.33 | 0 | 0 | 0.67 | 0 |
| *Leptodiscella* | 1.43 | 0 | 0 | 5.193 | 0 | 0.33 | 0 | 0 | 1 | 0 |
| *Lophiostoma* | 0 | 0 | 0 | 3.447 | 0 | 0 | 0 | 0 | 0.67 | 0 |
| *Pseudocoleophoma* | 1.2 | 1.39 | 1.317 | 3.323 | 1.203 | 0.33 | 0.33 | 0.33 | 1 | 0.33 |
| *Stachylidium* | 1.283 | 0.777 | 1.28 | 2.66 | 0 | 0.33 | 0.33 | 0.33 | 0.67 | 0 |
| ***D. uncinatum*** |  |  |  |  |  |  |  |  |  |  |
| *Alfaria* | 0 | 0 | 0 | 0.81 | 3.207 | 0 | 0 | 0 | 0.33 | 0.67 |
| *Bipolaris* | 0 | 1.407 | 0 | 0 | 2.157 | 0 | 0.33 | 0 | 0 | 0.67 |
| *Humicola* | 1.767 | 0 | 0 | 1.553 | 5.287 | 0.33 | 0 | 0 | 0.33 | 1 |
| *Scedosporium* | 1.667 | 0 | 0 | 0 | 3.437 | 0.33 | 0 | 0 | 0 | 0.67 |
| ***D. intortum*** |  |  |  |  |  |  |  |  |  |  |
| *Basidioascus* | 3.403 | 0 | 1.237 | 1.133 | 1.317 | 1 | 0 | 0.33 | 0.33 | 0.33 |
| *Coprinopsis* | 2.473 | 0 | 1.267 | 1.25 | 1.397 | 0.67 | 0 | 0.33 | 0.33 | 0.33 |
| *Cintractia* | 4.23 | 1.163 | 0 | 1.02 | 1.317 | 0.67 | 0.33 | 0 | 0.33 | 0.33 |

**Supplementary table 5:** Core soil fungal genera in *Desmodium* spp. plots

|  | Abundance |  |  |  |  | OCC |  |  |  |  |
| --- | --- | --- | --- | --- | --- | --- | --- | --- | --- | --- |
| Taxa | *D. incanum* | *D. intortum* | *D. ramosissimum* | *D. repandum* | *D. uncinatum* | *D. incanum* | *D. intortum* | *D. ramosissimum* | *D. repandum* | *D. uncinatum* |
| *Acrophialophora* | 5.49 | 4.757 | 6.757 | 5.287 | 6.283 | 0.67 | 0.67 | 1 | 0.67 | 1 |
| *Acrocalymma* | 6.81 | 7.013 | 6.59 | 5.067 | 4.137 | 1 | 1 | 1 | 1 | 0.67 |
| *Acremonium* | 7.217 | 6.587 | 7.41 | 6.187 | 7.233 | 1 | 1 | 1 | 1 | 1 |
| *Achroistachys* | 7.677 | 7.29 | 8.593 | 7.46 | 8.333 | 1 | 1 | 1 | 1 | 1 |
| *Alternaria* | 8.01 | 7.943 | 5.62 | 7.26 | 7.477 | 1 | 1 | 1 | 1 | 1 |
| *Auxarthron* | 6.653 | 2.8 | 7.147 | 6.637 | 5.607 | 1 | 0.67 | 1 | 1 | 1 |
| *Hannaella* | 8.793 | 9.61 | 7.223 | 7.197 | 9.42 | 1 | 1 | 1 | 1 | 1 |
| *Chaetomium* | 11.24 | 11.043 | 11.267 | 11.47 | 11.1 | 1 | 1 | 1 | 1 | 1 |
| *Cercospora* | 8.61 | 7.663 | 2.937 | 9.15 | 7.94 | 1 | 1 | 0.67 | 1 | 1 |
| *Cercophora* | 5.093 | 4.193 | 3.497 | 4.157 | 2.927 | 1 | 0.67 | 0.67 | 1 | 0.67 |
| *Ceratobasidium* | 5.28 | 5.187 | 5.807 | 4.303 | 5.483 | 0.67 | 0.67 | 1 | 0.67 | 1 |
| *Coniocessia* | 6.797 | 5.253 | 6.187 | 6.42 | 6.08 | 1 | 1 | 1 | 1 | 1 |
| *Colletotrichum* | 7.377 | 7.077 | 6.867 | 7.967 | 8.893 | 1 | 1 | 1 | 1 | 1 |
| *Clonostachys* | 5.303 | 7.7 | 6.09 | 6.637 | 6.57 | 1 | 1 | 1 | 1 | 1 |
| *Clitopilus* | 6.677 | 3.537 | 4.977 | 6.007 | 4.38 | 1 | 1 | 1 | 1 | 0.67 |
| *Cladorrhinum* | 10.203 | 9.113 | 9.76 | 9.103 | 9.95 | 1 | 1 | 1 | 1 | 1 |
| *Fusarium* | 12.41 | 12.4 | 12.677 | 12.367 | 12.51 | 1 | 1 | 1 | 1 | 1 |
| *Fusariella* | 4.65 | 4.17 | 4.263 | 5.14 | 4.85 | 0.67 | 0.67 | 1 | 1 | 1 |
| *Curvularia* | 8.443 | 9.413 | 9.153 | 8.093 | 9.56 | 1 | 1 | 1 | 1 | 1 |
| *Zopfiella* | 8.053 | 7.237 | 7.53 | 8.203 | 5.23 | 1 | 1 | 1 | 1 | 1 |
| *Westerdykella* | 2.783 | 2.837 | 4.343 | 4.553 | 4.057 | 0.67 | 0.67 | 1 | 1 | 1 |
| *Aspergillus* | 8.397 | 8.65 | 8.927 | 9.093 | 8.543 | 1 | 1 | 1 | 1 | 1 |
| *unidentified* | 11.223 | 12.163 | 11.077 | 10.993 | 11.963 | 1 | 1 | 1 | 1 | 1 |
| *Unclassified* | 13.243 | 13.347 | 13.433 | 13.427 | 13.343 | 1 | 1 | 1 | 1 | 1 |
| *Trichoderma* | 6.453 | 3.653 | 5.28 | 7.35 | 5.293 | 1 | 0.67 | 1 | 1 | 1 |
| *Torula* | 7.47 | 7.507 | 6.41 | 7.98 | 8.38 | 1 | 1 | 1 | 1 | 1 |
| *Thanatephorus* | 5.4 | 4.383 | 5.237 | 2.65 | 5.427 | 1 | 0.67 | 1 | 0.67 | 1 |
| *Tetracladium* | 5.29 | 4.397 | 5.76 | 3.077 | 5.297 | 1 | 1 | 1 | 0.67 | 1 |
| *Talaromyces* | 8.123 | 8.203 | 8.127 | 8.773 | 7.823 | 1 | 1 | 1 | 1 | 1 |
| *Stachybotrys* | 8.873 | 8.663 | 9.29 | 9.047 | 9.483 | 1 | 1 | 1 | 1 | 1 |
| *Didymella* | 11.513 | 11.857 | 11.423 | 11.21 | 11.937 | 1 | 1 | 1 | 1 | 1 |
| *Idriella* | 6.173 | 7.343 | 4.953 | 6.847 | 4.867 | 1 | 1 | 1 | 1 | 1 |
| *Scytalidium* | 2.867 | 3.267 | 3.45 | 4.98 | 4.857 | 0.67 | 0.67 | 1 | 1 | 1 |
| *Sclerostagonospora* | 6.433 | 5.563 | 6.033 | 4.26 | 2.93 | 1 | 1 | 1 | 0.67 | 0.67 |
| *Schizothecium* | 7.75 | 6.747 | 7.007 | 7.133 | 6.203 | 1 | 1 | 1 | 1 | 1 |
| *Lectera* | 6.803 | 4.467 | 8.783 | 5.543 | 6.347 | 1 | 1 | 1 | 1 | 1 |
| *Papiliotrema* | 8.117 | 5.773 | 7.68 | 5.44 | 6.88 | 1 | 1 | 1 | 1 | 1 |
| *Ochroconis* | 5.643 | 5.597 | 2.613 | 5.67 | 3.973 | 1 | 1 | 0.67 | 1 | 1 |
| *Nigrospora* | 7.843 | 7.717 | 6.977 | 6.463 | 7.037 | 1 | 1 | 1 | 1 | 1 |
| *Neurospora* | 4.72 | 6.52 | 5.123 | 5.317 | 3.083 | 0.67 | 1 | 1 | 1 | 0.67 |
| *Myrothecium* | 8.727 | 8.487 | 8.377 | 8.303 | 9.12 | 1 | 1 | 1 | 1 | 1 |
| *Mycosphaerella* | 9.53 | 9.87 | 7.107 | 6.773 | 9.087 | 1 | 1 | 1 | 1 | 1 |
| *Mortierella* | 8.087 | 8.487 | 8.187 | 6.55 | 8.117 | 1 | 1 | 1 | 1 | 1 |
| *Microdochium* | 6.34 | 4.86 | 5.637 | 5.32 | 4.483 | 1 | 0.67 | 1 | 1 | 1 |
| *Metarhizium* | 2.487 | 2.973 | 2.717 | 3.187 | 2.967 | 0.67 | 0.67 | 0.67 | 1 | 1 |
| *Sarocladium* | 2.847 | 2.827 | 4.097 | 3.143 | 2.783 | 0.67 | 0.67 | 1 | 0.67 | 0.67 |
| *Roussoella* | 4.16 | 1.8 | 4.513 | 3.24 | 3.65 | 1 | 0.67 | 1 | 1 | 0.67 |
| *Preussia* | 7.49 | 6.87 | 8.66 | 8.417 | 6.853 | 1 | 1 | 1 | 1 | 1 |
| *Plectosphaerella* | 7.757 | 7.833 | 9.143 | 10.33 | 8.673 | 1 | 1 | 1 | 1 | 1 |
| *Phaeosphaeria* | 9.237 | 9.123 | 8.973 | 7.823 | 9.417 | 1 | 1 | 1 | 1 | 1 |
| *Periconia* | 7.45 | 8.23 | 8.787 | 7.567 | 8.81 | 1 | 1 | 1 | 1 | 1 |
| *Penicillium* | 6.42 | 5.02 | 3.773 | 6.08 | 4.653 | 1 | 1 | 0.67 | 1 | 0.67 |
| *Paracremonium* | 7.017 | 6.397 | 6.817 | 7.197 | 6.603 | 1 | 1 | 1 | 1 | 1 |
| *Pyrenochaetopsis* | 6.607 | 8.16 | 4.22 | 5.697 | 9.577 | 1 | 1 | 0.67 | 1 | 1 |
| *Purpureocillium* | 4.237 | 3.407 | 4.653 | 5.9 | 5.603 | 1 | 0.67 | 1 | 1 | 1 |

**Supplementary table 6:** Pan fungal taxa shared among *Desmodium* spp. plots

| Taxa | *Desmodium* spp.<br>plots sharing taxa | Abundance |  |  |  |  | OCC |  |  |  |  |
| --- | --- | --- | --- | --- | --- | --- | --- | --- | --- | --- | --- |
|  |  | *D. incanum* | *D. intortum* | *D. ramosissimum* | *D. repandum* | *D. uncinatum* | *D. incanum* | *D. intortum* | *D. ramosissimum* | *D. repandum* | *D. uncinatum* |
| *Sporisorium* | DRM, DUN | 1.283 | 1.88 | 2.37 | 0 | 2.553 | 0.33 | 0.33 | 0.67 | 0 | 0.67 |
| *Stagonospora* | DRM, DUN | 1.21 | 1.877 | 3.24 | 0.637 | 4.307 | 0.33 | 0.33 | 0.67 | 0.33 | 0.67 |
| *Saitozyma* | DIN, DIT, DRM, DRP | 4.013 | 3.357 | 4.84 | 1.893 | 1.7 | 0.67 | 0.67 | 1 | 0.67 | 0.33 |
| *Setophaeosphaeria* | DIT, DRM, DUN | 2.06 | 4.107 | 2.447 | 5.357 | 3.783 | 0.33 | 0.67 | 0.33 | 1 | 0.67 |
| *Subulicystidium* | DRM, DRP | 0.877 | 0 | 1.247 | 1.747 | 0.717 | 0.33 | 0 | 0.67 | 0.67 | 0.33 |
| *Robillarda* | DIT, DRM, DUN | 1.43 | 6.347 | 1.677 | 5.923 | 8.303 | 0.33 | 1 | 0.33 | 1 | 1 |
| *Rhodosporiobolus* | DIN, DRM, DRP, DUN | 3.857 | 1.51 | 4.563 | 1.283 | 2.307 | 1 | 0.33 | 1 | 0.67 | 0.67 |
| *Rhizophlyctis* | DIN DIT DUN | 2.92 | 2.21 | 0 | 2.15 | 3.86 | 0.67 | 0.67 | 0 | 0.33 | 1 |
| *Pseudorobillarda* | DIN, DRM, DUN | 4.777 | 2.077 | 0.797 | 2.623 | 1.977 | 1 | 0.33 | 0.33 | 0.67 | 0.67 |
| *Psathyrella* | DIN, DRP | 2.933 | 1.843 | 0.593 | 2.563 | 0 | 0.67 | 0.33 | 0.33 | 0.67 | 0 |
| *Podospora* | DIN, DIT, DRP | 5.607 | 2.99 | 1.14 | 3.45 | 0 | 1 | 0.67 | 0.33 | 0.67 | 0 |
| *Poaceascoma* | DIN, DRM, DRP, DUN | 4.207 | 1.607 | 5.013 | 2.65 | 3.617 | 1 | 0.33 | 1 | 0.67 | 0.67 |
| *Phialophora* | DRP, DUN | 0.877 | 0 | 1.84 | 4.433 | 5.077 | 0.33 | 0 | 0.33 | 1 | 1 |
| *Montagnula* | DRP, DUN | 0 | 0 | 0.94 | 2.83 | 3.1 | 0 | 0 | 0.33 | 0.67 | 0.67 |
| *Monosporascus* | DIN, DIT, DRM, DUN | 2.307 | 3.147 | 1.75 | 0 | 4.593 | 0.67 | 0.67 | 0.67 | 0 | 0.67 |
| *Monographella* | DIN, DRP | 2.88 | 2.187 | 1.217 | 2.287 | 1.583 | 0.67 | 0.33 | 0.33 | 0.67 | 0.33 |
| *Modicella* | DRM, DUN | 2.71 | 0 | 6.063 | 0 | 3.71 | 0.33 | 0 | 1 | 0 | 0.67 |
| *Macrophomina* | DIN, DUN | 2.133 | 1.1 | 1.857 | 0 | 3.797 | 0.67 | 0.33 | 0.33 | 0 | 0.67 |
| *Metacordyceps* | DIT, DRM, DRP, DUN | 1.047 | 5.647 | 2.877 | 5.913 | 3.847 | 0.33 | 1 | 0.67 | 1 | 1 |
| *Magnaporthe* | DIN, DIT, DRM, DRP | 1.923 | 2.877 | 2.303 | 2.187 | 0 | 0.67 | 0.67 | 0.67 | 0.67 | 0 |
| *Microascus* | DIN, DIT, DRM | 5.227 | 2.263 | 3.483 | 0 | 1.363 | 1 | 0.67 | 1 | 0 | 0.33 |
| *Leucosphaerina* | DIT, DRP, DUN | 0.787 | 2.07 | 1.157 | 1.88 | 2.147 | 0.33 | 0.67 | 0.33 | 0.67 | 0.67 |
| *Leptospora* | DIT, DRP | 1.5 | 2.847 | 0 | 2.32 | 0 | 0.33 | 0.67 | 0 | 0.67 | 0 |
| *Kamienskia* | DIT, DUN | 1.423 | 4.737 | 1.237 | 0 | 4.053 | 0.33 | 1 | 0.33 | 0 | 1 |
| *Hirsutella* | DIT, DRP | 1.317 | 4.01 | 0 | 2.527 | 1.237 | 0.33 | 1 | 0 | 0.67 | 0.33 |
| *Hansfordia* | DIT, DRP, DUN | 1.2 | 4.433 | 0 | 4.943 | 3.627 | 0.33 | 1 | 0 | 1 | 1 |
| *Myrmecridium* | DIN, DIT, DRP, DUN | 3.267 | 3.807 | 1.45 | 2.777 | 4.107 | 0.67 | 0.67 | 0.33 | 0.67 | 1 |
| *Paracamarosporium* | DIN, DIT, DRM, DRP | 4.653 | 5.08 | 2.793 | 2.19 | 0 | 1 | 1 | 0.67 | 0.67 | 0 |
| *Xylaria* | DIN, DRP, DUN | 4.39 | 0 | 0 | 2.6 | 1.797 | 1 | 0 | 0 | 1 | 0.67 |
| *Conocybe* | DIT, DRM, DRP, DUN | 1.467 | 5.103 | 5.533 | 5.383 | 7.487 | 0.33 | 1 | 1 | 1 | 1 |
| *Cryptococcus* | DIN, DRM, DRP, DUN | 5.06 | 3.767 | 0.94 | 3.047 | 3.873 | 1 | 1 | 0.33 | 0.67 | 1 |
| *Dendryphiella* | DIT, DRM, DRP, DUN | 1.977 | 2.673 | 4.493 | 6.237 | 4.98 | 0.33 | 0.67 | 1 | 1 | 1 |
| *Chrysosporium* | DIN, DRM, DRP, DUN | 4.817 | 1.1 | 5.337 | 3.023 | 0 | 1 | 0.33 | 1 | 0.67 | 0 |
| *Chalara* | DIT, DRM, DUN | 1.767 | 4.12 | 2.24 | 1.403 | 2.71 | 0.33 | 0.67 | 0.67 | 0.33 | 0.67 |
| *Botryosphaeria* | DRM, DRP, DUN | 2.14 | 1.783 | 4.58 | 3.887 | 2.583 | 0.33 | 0.33 | 1 | 1 | 0.67 |
| *Boerlagiomyces* | DIN, DIT | 3.367 | 2.36 | 1.467 | 1.67 | 0 | 0.67 | 0.67 | 0.33 | 0.33 | 0 |
| *Bartalinia* | DRP, DUN | 0.95 | 1.14 | 0 | 1.917 | 3.01 | 0.33 | 0.33 | 0 | 0.67 | 0.67 |
| *Ascochyta* | DIN, DIT, DRM, DUN | 4.113 | 5.887 | 2.917 | 1.73 | 6.107 | 1 | 1 | 0.67 | 0.33 | 1 |
| *Arxiella* | DIT, DRP, DUN | 1.38 | 3.663 | 1.423 | 2.89 | 4.987 | 0.33 | 0.67 | 0.33 | 0.67 | 1 |
| *Arthrographis* | DRM, DRP | 2.147 | 1.42 | 2.133 | 2.997 | 0 | 0.33 | 0.33 | 0.67 | 0.67 | 0 |
| *Arthrobotrys* | DIN, DRP, DUN | 2.25 | 1.327 | 1.207 | 3.863 | 4.48 | 0.67 | 0.33 | 0.33 | 1 | 1 |
| *Arachnomycetes* | DIN, DRP, DUN | 2.977 | 0 | 1.267 | 2.1 | 2.637 | 0.67 | 0 | 0.33 | 0.67 | 0.67 |
| *Aplosporella* | DIT, DRM, DRP, DUN | 1.97 | 6.16 | 6.327 | 3.047 | 4.147 | 0.33 | 1 | 1 | 0.67 | 0.67 |
| *Antennariella* | DIN, DRM, DRP, DUN | 4.553 | 2.427 | 7.043 | 5.81 | 7.487 | 1 | 0.33 | 1 | 1 | 1 |
| *Wardomycopsis* | DIT, DRM, DRP, DUN | 1.747 | 2.523 | 5.187 | 3.877 | 3.383 | 0.33 | 0.67 | 1 | 1 | 0.67 |
| *Vishniacozyma* | DIT, DRM, DRP, DUN | 1.65 | 3.52 | 3.633 | 4.053 | 4.527 | 0.33 | 0.67 | 0.67 | 1 | 1 |
| *Funneliformis* | DIN, DIT, DUN | 3 | 2.743 | 0 | 0.743 | 3.857 | 0.67 | 0.67 | 0 | 0.33 | 0.67 |
| *Exserohilum* | DIN, DIT, DRM | 2.84 | 4.47 | 2.637 | 1.02 | 1.517 | 0.67 | 1 | 0.67 | 0.33 | 0.33 |
| *Entoloma* | DIN, DUN | 1.983 | 1.06 | 1.823 | 0 | 2.393 | 0.67 | 0.33 | 0.33 | 0 | 0.67 |
| *Endophragmiella* | DIN, DIT, DR, DRP | 3.85 | 4.993 | 5.493 | 5.787 | 1.483 | 1 | 1 | 1 | 1 | 0.33 |
| *Dioszegia* | DIN, DIT, DUN | 2.42 | 4.447 | 0 | 1.783 | 3.67 | 0.67 | 1 | 0 | 0.33 | 1 |
| *Graphium* | DRM, DRP, DUN | 0.98 | 1.163 | 4.62 | 4.45 | 4.457 | 0.33 | 0.33 | 1 | 1 | 1 |
| *Gibellulopsis* | DIN, DIT, DRP, DUN | 5.427 | 3.913 | 1.397 | 7.47 | 6.967 | 1 | 1 | 0.33 | 1 | 1 |
| *Lecanicillium* | DIT, DRM, DRP | 0.68 | 7.267 | 2.567 | 4.737 | 1.917 | 0.33 | 1 | 0.67 | 1 | 0.33 |
| *Lasiodiplodia* | DIN, DIT, DRM, DRP | 3.303 | 5.347 | 6.67 | 5.09 | 0.963 | 1 | 1 | 1 | 1 | 0.33 |

**Legend:** DRM = *D. ramosissimum*, DUN = *D. uncinatum*, DIN = *D. incanum*, DIT = *D. intortum*, DRP = *D. repandum*

## References

Barkai-Golan, R. (2008). Chapter 6—Aspergillus Mycotoxins. In R. Barkai-Golan & N. Paster (Eds.), Mycotoxins in Fruits and Vegetables (pp. 115–151). Academic Press. https://doi.org/10.1016/B978-0-12-374126-4.00006-1

Bokulich, N. A., Kaehler, B. D., Rideout, J. R., Dillon, M., Bolyen, E., Knight, R., Huttley, G. A., & Caporaso, J. G. (2018). Optimizing taxonomic classification of marker-gene amplicon sequences with QIIME 2’s q2-feature-classifier plugin. Microbiome, 6(1), 90. https://doi.org/10.1186/s40168-018-0470-z

Bolyen, E., Rideout, J. R., Dillon, M. R., Bokulich, N. A., Abnet, C. C., Al-Ghalith, G. A., Alexander, H., Alm, E. J., Arumugam, M., Asnicar, F., Bai, Y., Bisanz, J. E., Bittinger, K., Brejnrod, A., Brislawn, C. J., Brown, C. T., Callahan, B. J., Caraballo-Rodríguez, A. M., Chase, J., … Caporaso, J. G. (2019). Reproducible, interactive, scalable and extensible microbiome data science using QIIME 2. Nature Biotechnology, 37(8), 852–857. https://doi.org/10.1038/s41587-019-0209-9

Callahan, B. J., McMurdie, P. J., Rosen, M. J., Han, A. W., Johnson, A. J. A., & Holmes, S. P. (2016). DADA2: High-resolution sample inference from Illumina amplicon data. Nature Methods, 13(7), 581. https://doi.org/10.1038/nmeth.3869

Compant, S., Samad, A., Faist, H., & Sessitsch, A. (2019). A review on the plant microbiome: Ecology, functions, and emerging trends in microbial application. Journal of Advanced Research, 19, 29–37. https://doi.org/10.1016/j.jare.2019.03.004

DeSantis, T. Z., Hugenholtz, P., Larsen, N., Rojas, M., Brodie, E. L., Keller, K., Huber, T., Dalevi, D., Hu, P., & Andersen, G. L. (2006). Greengenes, a chimera-checked 16S rRNA gene database and workbench compatible with ARB. Applied and Environmental Microbiology, 72(7), 5069–5072. https://doi.org/10.1128/AEM.03006-05

Drinkwater, L. E., Midega, C. A. O., Awuor, R., Nyagol, D., & Khan, Z. R. (2021). Perennial legume intercrops provide multiple belowground ecosystem services in smallholder farming systems. Agriculture, Ecosystems & Environment, 320, 107566. https://doi.org/10.1016/j.agee.2021.107566

Eisenhauer, N., Milcu, A., Sabais, A. C. W., Bessler, H., Brenner, J., Engels, C., Klarner, B., Maraun, M., Partsch, S., Roscher, C., Schonert, F., Temperton, V. M., Thomisch, K., Weigelt, A., Weisser, W. W., & Scheu, S. (2011). Plant diversity surpasses plant functional groups and plant productivity as driver of soil biota in the long term. PLoS ONE, 6(1), e16055. https://doi.org/10.1371/journal.pone.0016055

Farid, B., Rafiou, M., Marcellin, A., Durand, D.-N., Nabede, A., Sylvestre, A., Haziz, S., Adolphe, A., Aly, S., & Lamine, B.-M. (2018). Ethnobotanical survey of three species of Desmodium genus (Desmodium ramosissimum, Desmodium gangeticum and Desmodium adscendens) used in traditional medicine, Benin. International Journal of Sciences, 4(12), 26–33. https://doi.org/10.18483/ijsci.1860

Friman, J., Pineda, A., Loon, J. J. A. van, & Dicke, M. (2021). Bidirectional plant-mediated interactions between rhizobacteria and shoot-feeding herbivorous insects: A community ecology perspective. Ecological Entomology, 46(1), 1–10. https://doi.org/10.1111/een.12966.

Goettel, M. S., Koike, M., Kim, J. J., Aiuchi, D., Shinya, R., & Brodeur, J. (2008). Potential of Lecanicillium spp. For management of insects, nematodes and plant diseases. Journal of Invertebrate Pathology, 98(3), 256–261. https://doi.org/10.1016/j.jip.2008.01.009

Gonzaga, L. L., Costa, L. E. O., Santos, T. T., Araújo, E. F., & Queiroz, M. V. (2015). Endophytic fungi from the genus Colletotrichum are abundant in the Phaseolus vulgaris and have high genetic diversity. Journal of Applied Microbiology, 118(2), 485–496. https://doi.org/10.1111/jam.12696

Grunseich, J. M., Thompson, M. N., Aguirre, N. M., & Helms, A. M. (2020). The role of plant-associated microbes in mediating host-plant selection by insect herbivores. Plants, 9(1), 1–23. https://doi.org/10.3390/plants9010006

Haichar, F. el Z., Santaella, C., Heulin, T., & Achouak, W. (2014). Root exudates mediated interactions belowground. Soil Biology and Biochemistry, 77, 69–80. https://doi.org/10.1016/j.soilbio.2014.06.017

Hamayun, M., Khan, S. A., Iqbal, I., Na, C.-I., Khan, A. L., Hwang, Y.-H., Lee, B.-H., & Lee, I.-J. (2009). Chrysosporium pseudomerdarium produces gibberellins and promotes plant growth. The Journal of Microbiology, 47(4), 425–430. https://doi.org/10.1007/s12275-009-0268-6

Heuzé, V., Tran, G., Eugène, M., & Bastianelli, D. (2015, October 7). Silverleaf desmodium (Desmodium uncinatum). Silverleaf Desmodium (Desmodium Uncinatum). http://www.feedipedia.org/node/299

Heuzé, V., Tran, G., & Hassoun, P. (2017). Greenleaf desmodium (Desmodium intortum). https://www.feedipedia.org/node/303

Hooper, A. M., Caulfield, J. C., Hao, B., Pickett, J. A., Midega, C. A. O., & Khan, Z. R. (2015). Isolation and identification of Desmodium root exudates from drought tolerant species used as intercrops against Striga hermonthica. Phytochemistry, 117, 380–387. https://doi.org/10.1016/j.phytochem.2015.06.026

Hooper, A. M., Tsanuo, M. K., Chamberlain, K., Tittcomb, K., Scholes, J., Hassanali, A., Khan, Z. R., & Pickett, J. A. (2010). Isoschaftoside, a C-glycosylflavonoid from Desmodium uncinatum root exudate, is an allelochemical against the development of Striga. Phytochemistry, 71(8), 904–908. https://doi.org/10.1016/j.phytochem.2010.02.015

Htwe, A. Z., Moh, S. M., Soe, K. M., Moe, K., & Yamakawa, T. (2019). Effects of biofertilizer produced from Bradyrhizobium and Streptomyces griseoflavus on plant growth, nodulation, nitrogen fixation, nutrient uptake, and seed yield of mung bean, cowpea, and soybean. Agronomy, 9(2), 77. https://doi.org/10.3390/agronomy9020077

Huang, X.-F., Zhou, D., Lapsansky, E. R., Reardon, K. F., Guo, J., Andales, M. J., Vivanco, J. M., & Manter, D. K. (2017). Mitsuaria sp. And Burkholderia sp. from Arabidopsis rhizosphere enhance drought tolerance in Arabidopsis thaliana and maize (Zea mays L.). Plant and Soil, 419(1), 523–539. https://doi.org/10.1007/s11104-017-3360-4

Jacoby, R., Peukert, M., Succurro, A., Koprivova, A., & Kopriva, S. (2017). The role of soil microorganisms in plant mineral nutrition—current knowledge and future directions. Frontiers in Plant Science, 8, 1617. https://doi.org/10.3389/fpls.2017.01617

Karunasekera, H., & Daniel, G. (2013). Molecular identification and phylogenic analysis by sequencing the rDNA of copper-tolerant soft-rot Phialophora spp. International Biodeterioration & Biodegradation, 82, 45–52. https://doi.org/10.1016/j.ibiod.2013.01.019

Liu, Z.-P., Wang, B.-J., Liu, Y.-H., & Liu, S.-J. (2005). Novosphingobium taihuense sp. Nov., a novel aromatic-compound-degrading bacterium isolated from Taihu Lake, China. International Journal of Systematic and Evolutionary Microbiology, 55(3), 1229–1232. https://doi.org/10.1099/ijs.0.63468-0

Lu, X., Taylor, A. E., Myrold, D. D., & Neufeld, J. D. (2020). Expanding perspectives of soil nitrification to include ammonia7oxidizing archaea and comammox bacteria. Soil Science Society of America Journal, 84(2), 287–302. https://doi.org/10.1002/saj2.20029

Ma, X., Zheng, C., Hu, C., Rahman, K., & Qin, L. (2011). The genus Desmodium (Fabaceae)-traditional uses in Chinese medicine, phytochemistry and pharmacology. Journal of Ethnopharmacology, 138(2), 314–332. https://doi.org/10.1016/j.jep.2011.09.053

Maxwell, O., Midega, C., Obonyo, M., & Zeyaur, R. (2017). Distribution of Aspergillus and Fusarium ear rot causative fungi in soils under push-pull and maize monocropping system in Western Kenya. African Journal of Microbiology Research, 11, 1411–1421. https://doi.org/10.5897/AJMR2017.8685

Midega, C. A. O., Bruce, T. J. A., Pickett, J. A., & Khan, Z. R. (2015). Ecological management of cereal stemborers in African smallholder agriculture through behavioural manipulation. Ecological Entomology, 40(Suppl 1), 70–81. https://doi.org/10.1111/een.12216

Midega, C. A. O., Pittchar, J. O., Pickett, J. A., Hailu, G. W., & Khan, Z. R. (2018). A climate-adapted push-pull system effectively controls fall armyworm, Spodoptera frugiperda (J E Smith), in maize in East Africa. Crop Protection, 105, 10–15. https://doi.org/10.1016/j.cropro.2017.11.003

Midega, C. A. O., Wasonga, C. J., Hooper, A. M., Pickett, J. A., & Khan, Z. R. (2017). Drought-tolerant Desmodium species effectively suppress parasitic striga weed and improve cereal grain yields in western Kenya. Crop Protection, 98, 94–101. https://doi.org/10.1016/j.cropro.2017.03.018

Mutyambai, D. M., Bass, E., Luttermoser, T., Poveda, K., Midega, C. A. O., Khan, Z. R., & Kessler, A. (2019). More than “push” and “pull”? plant-soil feedbacks of maize companion cropping increase chemical plant defenses against herbivores. Frontiers in Ecology and Evolution, 7, 217. https://doi.org/10.3389/fevo.2019.00217

Mwakilili, A. D., Mwaikono, K. S., Herrera, S. L., Midega, C. A. O., Magingo, F., Alsanius, B., Dekker, T., & Lyantagaye, S. L. (2021). Long-term maize-Desmodium intercropping shifts structure and composition of soil microbiome with stronger impact on fungal communities. Plant and Soil. https://doi.org/10.1007/s11104-021-05082-w

Nicoletti, R., & Becchimanzi, A. (2020). Endophytism of Lecanicillium and Akanthomyces. Agriculture, 10(6), 205. https://doi.org/10.3390/agriculture10060205

Nilsson, R. H., Larsson, K.-H., Taylor, A. F. S., Bengtsson-Palme, J., Jeppesen, T. S., Schigel, D., Kennedy, P., Picard, K., Glöckner, F. O., Tedersoo, L., Saar, I., Kõljalg, U., & Abarenkov, K. (2018). The UNITE database for molecular identification of fungi: Handling dark taxa and parallel taxonomic classifications. Nucleic Acids Research, 47(D1), D259–D264. https://doi.org/10.1093/nar/gky1022

Njeru, N. K., Midega, C. A. O., Muthomi, J. W., Wagacha, J. M., & Khan, Z. R. (2020). Impact of push–pull cropping system on pest management and occurrence of ear rots and mycotoxin contamination of maize in western Kenya. Plant Pathology, 69(9), 1644–1654. https://doi.org/10.1111/ppa.13259

Owuor, M. J., Midega, C. A. O., Obonyo, M., & Khan, Z. R. (2018). Impact of companion cropping on incidence and severity of maize ear rots and mycotoxins in Western Kenya. African Journal of Agricultural Research, 41(13), 2224–2231. https://doi.org/10.5897/AJAR2018.13396

Pangesti, N., Pineda, A., Dicke, M., & Loon, J. J. A. van. (2015). Variation in plant-mediated interactions between rhizobacteria and caterpillars: Potential role of soil composition. Plant Biology (Stuttgart, Germany), 17(2), 474–483. https://doi.org/10.1111/plb.12265

Parker, M. A. (2002). Bradyrhizobia from wild phaseolus, Desmodium, and Macroptilium species in Northern Mexico. Applied and Environmental Microbiology, 68(4), 2044–2048. https://doi.org/10.1128/AEM.68.4.2044

Philippot, L., Raaijmakers, J. M., Lemanceau, P., & van der Putten, W. H. (2013). Going back to the roots: The microbial ecology of the rhizosphere. Nature Reviews Microbiology, 11(11), 789–799. https://doi.org/10.1038/nrmicro3109

Pickett, J. A., Woodcock, C. M., Midega, C. A. O., & Khan, Z. R. (2014). Push–pull farming systems. Current Opinion in Biotechnology, 26, 125–132. https://doi.org/10.1016/j.copbio.2013.12.006

Rivas, R., Trujillo, M. E., Mateos, P. F., Martínez-Molina, E., & Velázquez, E. (2004). Agromyces ulmi sp. Nov., a xylanolytic bacterium isolated from Ulmus nigra in Spain. International Journal of Systematic and Evolutionary Microbiology, 54(Pt 6), 1987–1990. https://doi.org/10.1099/ijs.0.63058-0

Saccá, M. L., Barra Caracciolo, A., Di Lenola, M., & Grenni, P. (2017). Ecosystem services provided by soil microorganisms. In M. Lukac, P. Grenni, & M. Gamboni (Eds.), Soil Biological Communities and Ecosystem Resilience (pp. 9–24). Springer International Publishing.

Sarrocco, S., Mauro, A., & Battilani, P. (2019). Use of Competitive filamentous fungi as an alternative approach for mycotoxin risk reduction in staple cereals: state of art and future perspectives. Toxins, 11(12), 701. https://doi.org/10.3390/toxins11120701

Skiada, V., Avramidou, M., Bonfante, P., Genre, A., & Papadopoulou, K. K. (2020). An endophytic Fusarium–legume association is partially dependent on the common symbiotic signalling pathway. New Phytologist, 226(5), 1429–1444. https://doi.org/10.1111/nph.16457

Sohn, J. H., Kwon, K. K., Kang, J.-H., Jung, H.-B., & Kim, S.-J. (2004). Novosphingobium pentaromativorans sp. Nov., a high-molecular-mass polycyclic aromatic hydrocarbon-degrading bacterium isolated from estuarine sediment. International Journal of Systematic and Evolutionary Microbiology, 54(5), 1483–1487. https://doi.org/10.1099/ijs.0.02945-0

Summerell, B. A. (2019). Resolving Fusarium: current status of the genus. Annual Review of Phytopathology, 57(1), 323–339. https://doi.org/10.1146/annurev-phyto-082718-100204

Tivoli, B., & Banniza, S. (2007). Comparison of the epidemiology of ascochyta blights on grain legumes. European Journal of Plant Pathology, 119(1), 59–76. https://doi.org/10.1007/s10658-007-9117-9

Toniutti, M. A., Fornasero, L. V., Albicoro, F. J., Martini, M. C., Draghi, W., Alvarez, F., Lagares, A., Pensiero, J. F., & Papa, M. F. D. (2017). Nitrogen-fixing rhizobial strains isolated from Desmodium incanum DC in Argentina: Phylogeny, biodiversity and symbiotic ability. Systematic and Applied Microbiology, 40(5), 297–307. https://doi.org/10.1016/j.syapm.2017.04.004

Vukicevich, E., Lowery, T., Bowen, P., Úrbez-Torres, J. R., & Hart, M. (2016). Cover crops to increase soil microbial diversity and mitigate decline in perennial agriculture. A review. Agronomy for Sustainable Development, 36(3), 48. https://doi.org/10.1007/s13593-016-0385-7

Vurukonda, S. S. K. P., Vardharajula, S., Shrivastava, M., & SkZ, A. (2016). Enhancement of drought stress tolerance in crops by plant growth promoting rhizobacteria. Microbiological Research, 184, 13–24. https://doi.org/10.1016/j.micres.2015.12.003

Wingett, S. W., & Andrews, S. (2018). FastQ Screen: A tool for multi-genome mapping and quality control. F1000Research, 7, 1338. https://doi.org/10.12688/f1000research.15931.2

Wu, Z., Liu, Q., Li, Z., Cheng, W., Sun, J., Guo, Z., Li, Y., Zhou, J., Meng, D., Li, H., Lei, P., & Yin, H. (2018). Environmental factors shaping the diversity of bacterial communities that promote rice production. BMC Microbiology, 18(1), 51. https://doi.org/10.1186/s12866-018-1174-z

Xiao, X., Chen, W., Zong, L., Yang, J., Jiao, S., Lin, Y., Wang, E., & Wei, G. (2017). Two cultivated legume plants reveal the enrichment process of the microbiome in the rhizocompartments. Molecular Ecology, 26(6), 1641–1651. https://doi.org/10.1111/mec.14027

Xu, K. W., Zou, L., Penttinen, P., Zeng, X., Liu, M., Zhao, K., Chen, C., Chen, Y. X., & Zhang, X. (2016). Diversity and phylogeny of rhizobia associated with Desmodium spp. in Panxi, Sichuan, China. Systematic and Applied Microbiology, 39(1), 33–40. https://doi.org/10.1016/j.syapm.2015.10.005

Xu, L., Ravnskov, S., Larsen, J., Nilsson, R. H., & Nicolaisen, M. (2012). Soil fungal community structure along a soil health gradient in pea fields examined using deep amplicon sequencing. Soil Biology and Biochemistry, 46, 26–32. https://doi.org/10.1016/j.soilbio.2011.11.010

Zakrzewski, M., Proietti, C., Ellis, J. J., Hasan, S., Brion, M.-J., Berger, B., & Krause, L. (2017). Calypso: A user-friendly web-server for mining and visualizing microbiome-environment interactions. Bioinformatics (Oxford, England), 33(5), 782–783. https://doi.org/10.1093/bioinformatics/btw725

Zolla, G., Badri, D. V., Bakker, M. G., Manter, D. K., & Vivanco, J. M. (2013). Soil microbiomes vary in their ability to confer drought tolerance to Arabidopsis. Applied Soil Ecology, 68, 1–9. https://doi.org/10.1016/j.apsoil.2013.03.007

Zriba, N., Sherwood, J. E., & Mathre, D. E. (1999). Characterization and effectiveness of Phialophora spp. isolated from a Montana take-all suppressive soil in controlling take-all disease of wheat. Canadian Journal of Plant Pathology, 21(2), 110–118. https://doi.org/10.1080/07060669909501200

